# Cortical asymmetry and its developmental change have distinct relationships with brain connectivity

**DOI:** 10.64898/2026.10.28.762579

**Authors:** Siva Venkadesh, Fang-Cheng Yeh, Patrick R. Hof, Julie R. Korenberg

**Affiliations:** Department of Neurological Surgery, University of Pittsburgh, Pittsburgh, PA, USA; Nash Family Department of Neuroscience and Friedman Brain Institute, Icahn School of Medicine at Mount Sinai, New York, NY, USA; Departments of Pediatrics and Neurology, University of Utah, Salt Lake City, UT, USA

**Author notes:** Co-senior authors (Patrick R. Hof, Julie R. Korenberg).

**Keywords:** hemispheric asymmetry, corpus callosum, brain development, structural connectivity, tractography

## Abstract

The two cerebral hemispheres specialize while remaining interconnected. Does greater structural asymmetry imply less connectivity, and does connectivity also relate to how asymmetry changes during development? We addressed these questions in three independent cohorts spanning birth to 22 years, measuring asymmetry as the magnitude of the left–right difference in association tract overlap across 74 homotopic cortical region pairs. The regional pattern of asymmetry was similar across cohorts. In all three cohorts, more asymmetric regions had lower connectional density to cortical and non-cortical targets. However, involvement of the corpus callosum, the major pathway connecting the hemispheres, showed no consistent inverse relationship with regional asymmetry across cohorts. Instead, callosal involvement conditioned the relationship between mean regional asymmetry and its age-related change. In the two older cohorts, greater mean asymmetry accompanied more positive age effects among regions with appreciable callosal overlap, whereas this relationship was weak or negative among regions with little callosal overlap (standardized interactions +0.42 and +0.56). The pattern replicated across the two cohorts at matched ages, and longitudinal region-mean estimates in the Adolescent Brain Cognitive Development cohort were similar in direction and magnitude. In the youngest cohort, regions converged toward the mean, narrowing the between-region spread by 12%, with no detected callosal moderation. More asymmetric regions have lower overall connectional density, whereas callosal involvement helps explain how regional asymmetry changes with age.

**Significance statement:** The brain’s hemispheres differ in structure and function while remaining extensively connected. Does connectivity relate to how asymmetric a region is, how its asymmetry develops, or both? We distinguish these questions in three developmental cohorts. More asymmetric regions had lower overall connectional density. In contrast, involvement of the corpus callosum, the major pathway linking the hemispheres, showed no consistent inverse relationship with asymmetry across cohorts. Instead, it distinguished how mean regional asymmetry related to age-related change in the two older cohorts. A static comparison would miss this developmental relationship. These findings provide a framework for studying hemispheric development and investigating whether connectivity–asymmetry relationships are altered in neurodevelopmental conditions.

## Introduction

The cerebral hemispheres differ in their anatomy and functional specialization, yet communicate through extensive connections, principally via the corpus callosum. How do these differences coexist with connectivity? Hemispheric asymmetry is a widespread organizing feature of nervous systems (1), and human cortical and white-matter asymmetries are already evident in infancy (2–4). Their early presence raises a developmental question: does a region’s connectivity relate only to how asymmetric it is, or also to how that asymmetry changes as the brain develops?

One influential account links hemispheric specialization to the time required for signals to travel between hemispheres (5). Comparative and human allometric studies show that brain size relates to the balance of interhemispheric connectivity and specialization (6–8). Within humans, functional lateralization has been related to preferential communication within a hemisphere, callosal connectivity, and broader cortical organization (9–11). Callosal connections can also support both integration and interhemispheric inhibition (12, 13). Longitudinal work in the Adolescent Brain Cognitive Development study (ABCD) has linked earlier callosal microstructure to subsequent left–right differences in cortical thickness, cortical surface area and subcortical volume (14). Together, these studies link functional specialization and morphometric asymmetry to interhemispheric communication. Across cortical regions, it remains unclear how asymmetries in their overlap with association tracts relate to overall regional connectional density and local callosal involvement, and whether these relationships distinguish regional patterns of asymmetry development.

We separate three questions. Where does an asymmetric region sit in the broader network of brain connections? How much callosal involvement does it have? And does callosal involvement distinguish how its asymmetry changes with age? We measure structural asymmetry directly from the left–right difference in a region’s association tract overlap. This approach complements studies of functional lateralization and of asymmetries in particular white-matter pathways (15, 16). It also accommodates the possibility that different tracts within one region favor opposite hemispheres.

Three independent cohorts span birth to 22 years. The Baby Connectome Project (BCP) samples infancy and early childhood; the ABCD study provides three waves of repeated measurements; and HCP-Development (HCP-D) provides the widest age range. HCP-D’s age range encompasses ABCD’s, so the cohorts are complementary samples rather than successive developmental stages. We compare regional asymmetry with independently constructed region-to-region connectivity, then examine callosal involvement and age-related change, in all three cohorts. ABCD’s repeated scans additionally allow comparison with longitudinal change measured within individuals.

The results distinguish two relationships. More asymmetric regions have lower overall connectional density, but not consistently less callosal involvement. Callosal involvement instead helps explain how the relationship between mean regional asymmetry and age-related change differs across the cortex.

## Results

### A reproducible regional pattern of tract asymmetry

To relate cortical asymmetry to connectivity, we first needed a regional measure that preserves differences carried by distinct tract families, each a named association pathway (Methods). A cortical region can intersect several association pathways, and those pathways need not favor the same hemisphere. We therefore measured the absolute left–right difference in overlap for each tract family before averaging across the families covering a region. We then adjusted for total association-tract overlap, giving a regional measure of asymmetry beyond that expected from its overall overlap magnitude. Positive and negative adjusted values indicate greater and lesser asymmetry than expected, respectively; they do not indicate leftward and rightward dominance.

The planum temporale illustrates why tract families must be kept separate before taking their average (Fig. 1A). In HCP-D, arcuate fasciculus overlap was greater on the left (median 0.165 versus 0.016), whereas middle longitudinal fasciculus overlap was greater on the right (0.058 versus 0.150). The arcuate pattern is consistent with an earlier description of leftward asymmetry in this pathway (17).

**Figure 1.**
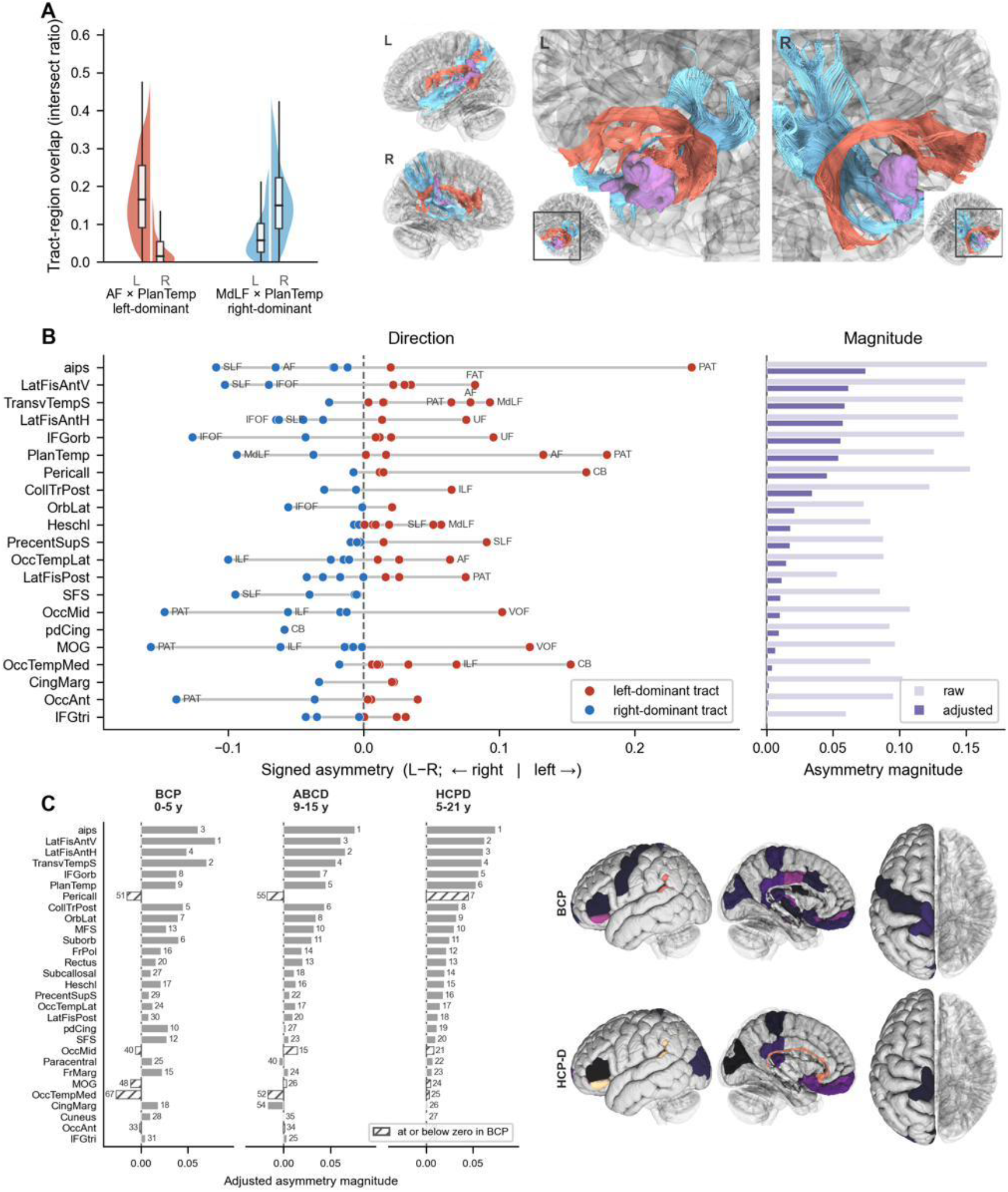
A reproducible regional pattern of tract asymmetry. (A) Left and right tract–region overlap in HCP-D for the arcuate fasciculus (AF; n = 630) and middle longitudinal fasciculus (MdLF; n = 633) at planum temporale, with tract renderings from an example participant. The two families favor opposite hemispheres. Each half-violin is a kernel density estimate independently scaled to equal maximum width. Lines and boxes show medians and interquartile ranges; whiskers extend to the most extreme observed values within 1.5 interquartile ranges beyond the quartiles. Values beyond the whiskers contribute to the density but are not plotted individually. (B) Signed family-level differences in HCP-D (left) and regional raw and adjusted magnitudes (right) for 21 regions with raw mean absolute difference ≥0.05 and positive adjusted asymmetry. Points represent families; horizontal lines span their range. Families with absolute group-mean differences >0.05 are labeled. (C) The 29 regions with positive adjusted asymmetry in HCP-D, shown in the same order across cohorts; numbers give within-cohort ranks among 74 regions. Hatching identifies regions at or below zero in BCP. Surface views show BCP and HCP-D. Adjusted asymmetry is the within-session residual on log total association overlap, averaged across sessions. Zero therefore marks the expected magnitude given total overlap, not anatomical symmetry. Samples: BCP, 450 sessions; ABCD, 19,569 sessions; HCP-D, 636 participants. Tract abbreviations and selected regional labels are defined in the SI Appendix, Table S10; aips denotes the anterior intermediate parietal sulcus.

Averaging signed differences would allow these opposing directions to cancel, understating the amount of asymmetry present within the region.

The underlying tract differences were consistent across participants. For the tract family with the highest sign consistency in each region, the median proportion agreeing on the sign of the left–right difference was 0.911; all 74 regions exceeded chance after Bonferroni correction across all 332 tested family–region cells. Additional tract-by-region validation, including opposing tract directions at Brodmann area 44, is provided in the SI Appendix, section S3. These checks establish reproducible directional differences at the tract level, while the regional measure summarizes their magnitude.

The regional pattern was similar across all three cohorts (Fig. 1B–C). Perisylvian, temporal and inferior parietal regions occupied the highest ranks, including the anterior intermediate parietal sulcus (sulcus intermedius primus of Jensen), the anterior rami of the lateral fissure, the transverse temporal sulcus, and planum temporale. Regional ordering was also stable when a common overlap-adjustment slope replaced the cohort-specific slopes (all ρ ≥ 0.992; SI Appendix, section S3 and Table S2). Differences in individual regional ranks are reported there as between-sample observations. Thus, a reproducible pattern of regional asymmetry provides the basis for asking how asymmetry relates to connectivity, without assigning each region a single dominant hemisphere.

### Connectional density and callosal involvement relate differently to asymmetry

Does greater regional asymmetry accompany lower connectivity, and does that relationship extend to connections through the corpus callosum? We first examined overall connectional density using a region-to-region graph reconstructed independently of the named-tract representation used to measure asymmetry, built with one pipeline in all three cohorts. Cortical degree summarizes a region’s mean edge weight to cortical targets in both hemispheres. Callosal involvement instead measures how much the region overlaps reconstructed callosal pathways. The two measures therefore describe different aspects of connectivity.

In all three cohorts, more asymmetric regions had lower cortical degree (ρ = −0.607 in HCP-D, −0.720 in BCP, and −0.559 in ABCD across 74 regions; all spin P < 0.001; Fig. 2A and SI Appendix, Table S3). The same pattern appeared within individual observations: the median correlation was −0.300, −0.312, and −0.302, and 99.7%, 99.3%, and 99.7% of the 636 HCP-D participants, 446 BCP sessions, and 1,000 ABCD participants showed a negative relationship. Lower density also extended to non-cortical targets, including subcortical and cerebellar gray matter (ρ = −0.496, −0.601, and −0.374 across regions). In HCP-D the cortical correlation was the larger of the two at full target breadth, but matching the number of cortical and non-cortical targets yielded correlations of similar magnitude (SI Appendix, section S4).

**Figure 2.**
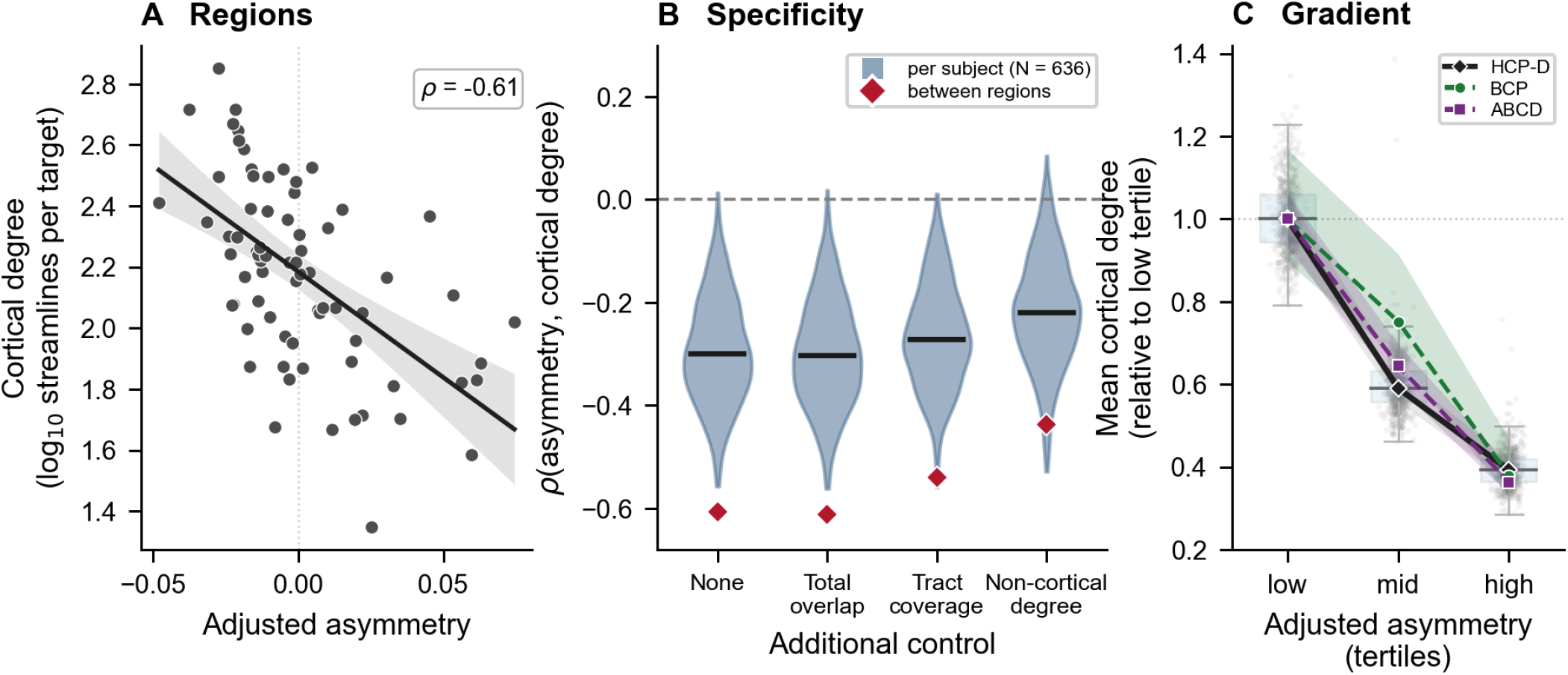
Greater regional asymmetry accompanies lower connectional density in three cohorts. 74 homotopic region pairs; HCP-D (636 participants), BCP (446 sessions), and ABCD (1,000 participants). Degree is computed from whole-brain tractography built with one pipeline in all cohorts (Methods). (A) HCP-D cortical degree, the mean number of streamlines connecting a region to cortical targets in both hemispheres, versus adjusted asymmetry. Points are regional means; the line and shading show an ordinary least-squares fit and 95% band on the log10 degree scale. The reported correlation is Spearman ρ. (B) HCP-D participant-level correlations (kernel density violins, with median bars) and between-region correlations (diamonds), before additional control and after separately controlling total association overlap, tract-family coverage, or non-cortical degree. These controls share variance with cortical degree and represent different components of the relationship. Values for all three cohorts are in SI Appendix, Table S3. (C) Mean cortical degree for each participant or BCP session within regional asymmetry tertiles (24–25 regions each, formed within each cohort), expressed relative to each cohort’s low-tertile median. Lines connect tertile medians; shaded bands span the interquartile range across participants or BCP sessions. HCP-D additionally shows boxes (interquartile range, with median lines) and all participants as points; a small number of HCP-D points above 1.4 lie outside the axis. Cortical and non-cortical degree use separate target sets (Methods).

The inverse cortical relationship persisted in every cohort after separately accounting for total tract overlap or non-cortical degree, and in BCP and HCP-D after accounting for the number of covered tract families (Fig. 2B; SI Appendix, Table S3). All 24 HCP-D regions in the high-asymmetry, low-degree group also showed sign consistency exceeding chance after correction across all tested family–region cells. Degree declined across asymmetry tertiles in all three cohorts, with high-tertile medians at 36– 39% of their respective low-tertile medians (Fig. 2C). These analyses support the inverse relationship across cohorts, participants, and measurement choices, while leaving shared tractography biases as a possible limitation.

Callosal involvement showed a different pattern (Fig. 3). Its regional correlation with adjusted asymmetry was +0.298 in BCP, −0.128 in ABCD, and −0.026 in HCP-D. The BCP and HCP-D correlations did not exceed their spatial nulls (spin P = 0.053 and 0.859). The most asymmetric tertile also did not have appreciably lower mean callosal involvement than the least asymmetric tertile in either displayed cohort (BCP, 0.155 versus 0.051; HCP-D, 0.114 versus 0.118). These observations do not establish the absence of all callosal relationships, but they provide no consistent inverse relationship with asymmetry magnitude.

**Figure 3.**
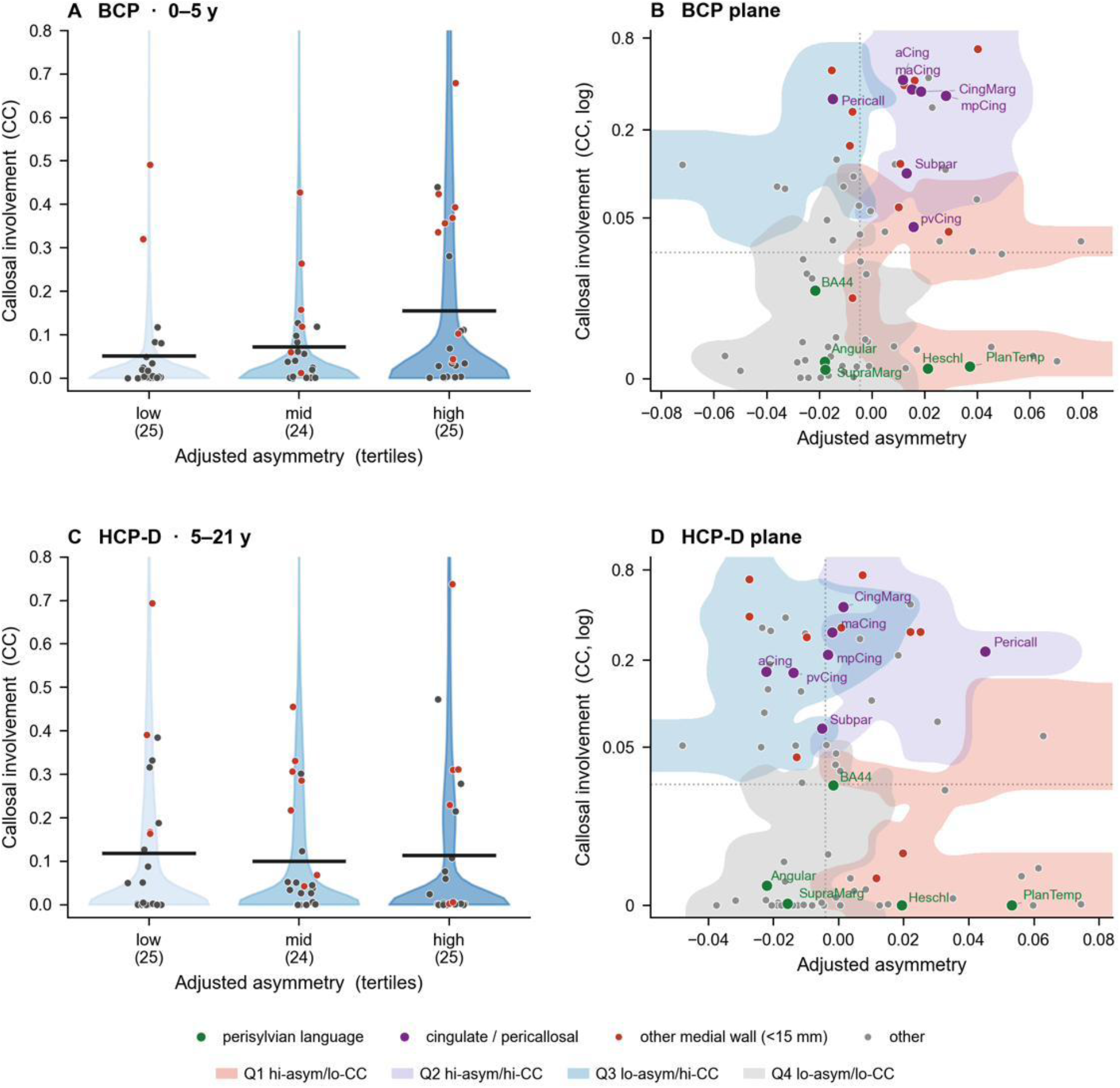
Callosal involvement has no consistent inverse relationship with asymmetry magnitude. BCP (450 sessions) and HCP-D (636 participants), each with 74 homotopic region pairs. (A, C) Callosal overlap within regional asymmetry tertiles. Violins show pooled session-by-region distributions, points show regional means, and horizontal bars show the mean across regions. Tertile sizes are given below the axes. (B, D) Regional adjusted asymmetry against unadjusted callosal overlap on a log scale. Shaded hulls display four descriptive combinations of higher or lower asymmetry and callosal involvement; their boundaries are not used for inference. Green labels identify perisylvian language regions and purple labels cingulate or pericallosal regions. Other regions within 15 mm of the midline are red; remaining regions are gray. Per-region sample sizes vary with coverage (BCP, 207– 450; HCP-D, 278–636). The ABCD correlation is reported in the SI Appendix, section S4 and Table S4. BA44: opercular part of the inferior frontal gyrus.

Greater asymmetry therefore accompanies lower overall connectional density without a corresponding, consistent reduction in callosal involvement. This distinction motivates the developmental question: does callosal involvement relate to how regional asymmetry changes with age?

### Callosal involvement distinguishes developmental patterns of asymmetry

A region that is highly asymmetric on average could show increasing, decreasing, or stable asymmetry with age. We asked whether callosal involvement helps distinguish these possibilities across regions. For each region, we estimated its age effect as the within-cohort correlation between age and adjusted asymmetry (ρ_age_), then related this age effect to its mean asymmetry and callosal involvement. Mean asymmetry is the average across the cohort, rather than an initial measurement.

In BCP, regions with greater mean asymmetry had more negative age effects (ρ = −0.413, spin P = 0.0022; Fig. 4A,C). This pattern persisted when mean asymmetry and age effects were estimated from disjoint groups of participants (ρ = −0.352; all 200 replications negative). The regional distribution also narrowed directly: sampling error-corrected between-region standard deviation declined by 12.1% across sample-size-matched age bins. Together, these results indicate convergence across infancy and early childhood: more asymmetric regions tended toward lower asymmetry with age, and less asymmetric regions toward higher asymmetry. The corresponding overall relationship was near zero in ABCD and positive but not established against the spatial null in HCP-D (SI Appendix, section S5 and Table S5).

**Figure 4.**
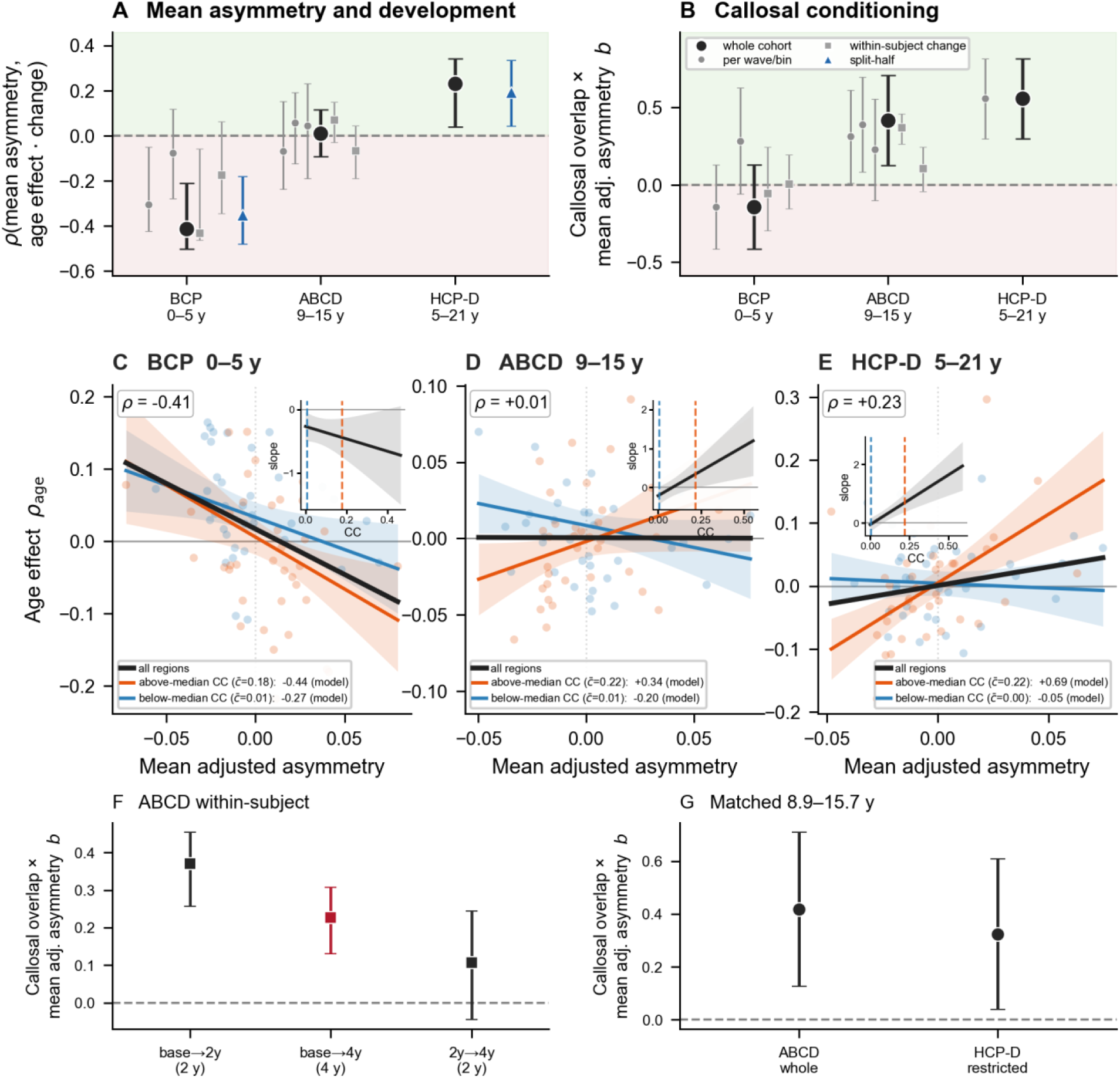
Callosal involvement distinguishes regional patterns of developmental change. BCP (429 sessions with usable age), ABCD (19,569 sessions from 10,652 participants), and HCP-D (636 participants); 74 regions per cohort. Mean asymmetry denotes the cohort mean; ρ_age_ = Spearman ρ(age, adjusted asymmetry). (A) Regional correlations of mean asymmetry with age effects (circles/triangles) or mean within-participant change (squares). (B) Standardized callosal-by-mean-asymmetry interaction. Large/small circles denote whole cohorts/waves or age bins; squares denote within-subject change. In (A), bars are bootstrap 95% percentile intervals resampling sessions or subject pairs. Blue triangles in BCP and HCP-D show the median and 2.5th– 97.5th percentiles across 200 disjoint half-sample replicates, describing replicate spread rather than full-sample confidence intervals. In (B), circles use regional ordinary least-squares 95% t intervals (74 regions; 70 degrees of freedom); squares use bootstrap 95% percentile intervals over subject pairs. Green/pink backgrounds mark positive/negative values, not uncertainty. (C–E) Age effects plotted against mean adjusted asymmetry. A positive interaction means that the relationship between mean adjusted asymmetry and the age effect becomes more positive as callosal overlap increases. Black lines are bivariate fits without bands. Colored lines evaluate the interaction model at the below- and above-median groups’ mean callosal overlap; model-implied slopes are labeled. Colored shading shows 95% confidence bands (±1.96 SE) for the fitted lines, calculated from model covariance, not prediction intervals. Insets show the fitted slope with 95% bands. Separate subgroup fits: SI Appendix, Table S5. (F) ABCD visit-transition interactions, with bootstrap 95% percentile intervals over subject pairs (2,000 replicates). (G) ABCD and HCP-D interactions at matched ages (8.9–15.7 years), with the same regional t intervals as B. Cohorts overlap in age and are not successive stages. Sensitivity analyses and interval comparisons are in the SI Appendix, sections S5 and S6.

In the two older cohorts, callosal involvement distinguished these patterns (Fig. 4B,D–E). Where callosal overlap was appreciable, greater average asymmetry went with a stronger tendency toward increasing asymmetry with age. Where overlap was low, that relationship was weak or reversed. This describes how age effects vary across regions; it does not mean that every region with appreciable overlap increased. The pattern was captured by positive standardized interactions between mean asymmetry and callosal involvement: +0.418 in ABCD and +0.558 in HCP-D. BCP’s interaction was −0.142, with no detected moderation.

The fitted lines in Fig. 4 show the interaction at representative callosal values, and the insets show how the fitted relationship varies continuously with callosal overlap. Because callosal overlap is strongly skewed, the comparison is largely between near-zero overlap and appreciable involvement. Callosal involvement alone did not explain the pattern consistently: its main effect was near zero in HCP-D, where the interaction was largest (SI Appendix, section S5).

The positive interactions persisted across the specification and influence checks in both older cohorts (SI Appendix, Table S6). Spatial checks gave spin P = 0.028 in ABCD and 0.0008 in HCP-D; their outcome-rotation null breaks the spatial relationship with all fixed model predictors, so these values assess spatial specificity rather than independently testing moderation. Adjustment for distance from the midline also retained positive interactions, as did excluding medial-wall regions. These checks address sensitivity to spatial position, although midline distance cannot separate callosal anatomy from distance-related imaging effects.

The pattern also replicated across the two older cohorts at matched ages (Fig. 4G): the interaction was +0.418 in ABCD and +0.325 in HCP-D restricted to the same 8.9–15.7-year window. Sex-stratified analyses detected no difference in the interaction (SI Appendix, section S8). Together, these results identify a callosal relationship with the regional pattern of developmental change in both older cohorts. The difference from BCP does not locate its developmental onset or show that it strengthens with age.

### Longitudinal measurements show a similar regional relationship

Do age differences among participants reflect a relationship also visible when the same individuals are followed over time? We addressed this question with ABCD’s repeated scans, calculating each region’s mean change between consecutive visits and fitting the same interaction model. The predictor remained cohort-mean regional asymmetry, rather than each participant’s earlier value, to avoid directly coupling individual change to its starting measurement. These analyses concern regional mean change, not prediction of individual trajectories.

Across 8,512 wave-adjacent visit pairs, the standardized interaction was +0.316, close to the mean of ABCD’s per-wave cross-sectional estimates (+0.311). Pooling consecutive transitions holds the nominal interval at two years; a sensitivity analysis including four-year pairs also gave a positive estimate (SI Appendix, section S6 and Table S7). The agreement in direction and magnitude supports continuity between the cross-sectional and longitudinal regional patterns.

The strength of the longitudinal inference was more limited. Its P value was 0.044 under ordinary least squares and 0.092 under the spatial null; the BCa bootstrap interval was [−0.023, +0.806]. Precision weighting gave stronger evidence, partly sensitive to model curvature (SI Appendix, section S6). The longitudinal estimate therefore provides agreement in magnitude, with uncertainty that depends on the estimator.

The three visit transitions also differed (Fig. 4F). The baseline-to-two-year interaction was larger than the baseline-to-four-year estimate, and the two consecutive two-year estimates differed numerically. The reported checks did not explain these differences, so they are not interpreted as an age trend.

Individual change was much noisier than region-mean change, with a shared-wave noise model attributing 97.4% of individual-change variance to measurement error under its assumptions (SI Appendix, section S6). Longitudinal measurements thus recover a similar regional relationship, while providing limited inferential confirmation and little basis for individual prediction.

### The regional pattern of change differs between BCP and the older cohorts

Which regions contribute to these developmental patterns, and are they the same across cohorts? The age-effect maps agreed moderately between ABCD and HCP-D (observed ρ = +0.53; +0.657 after correcting for map reliability), whereas BCP’s map showed little agreement with either older cohort (Fig. 5; SI Appendix, section S7 and Table S8). Change maps were less reliable than maps of mean asymmetry, particularly in BCP and HCP-D, and the map comparisons are descriptive rather than spatially tested replications.

**Figure 5.**
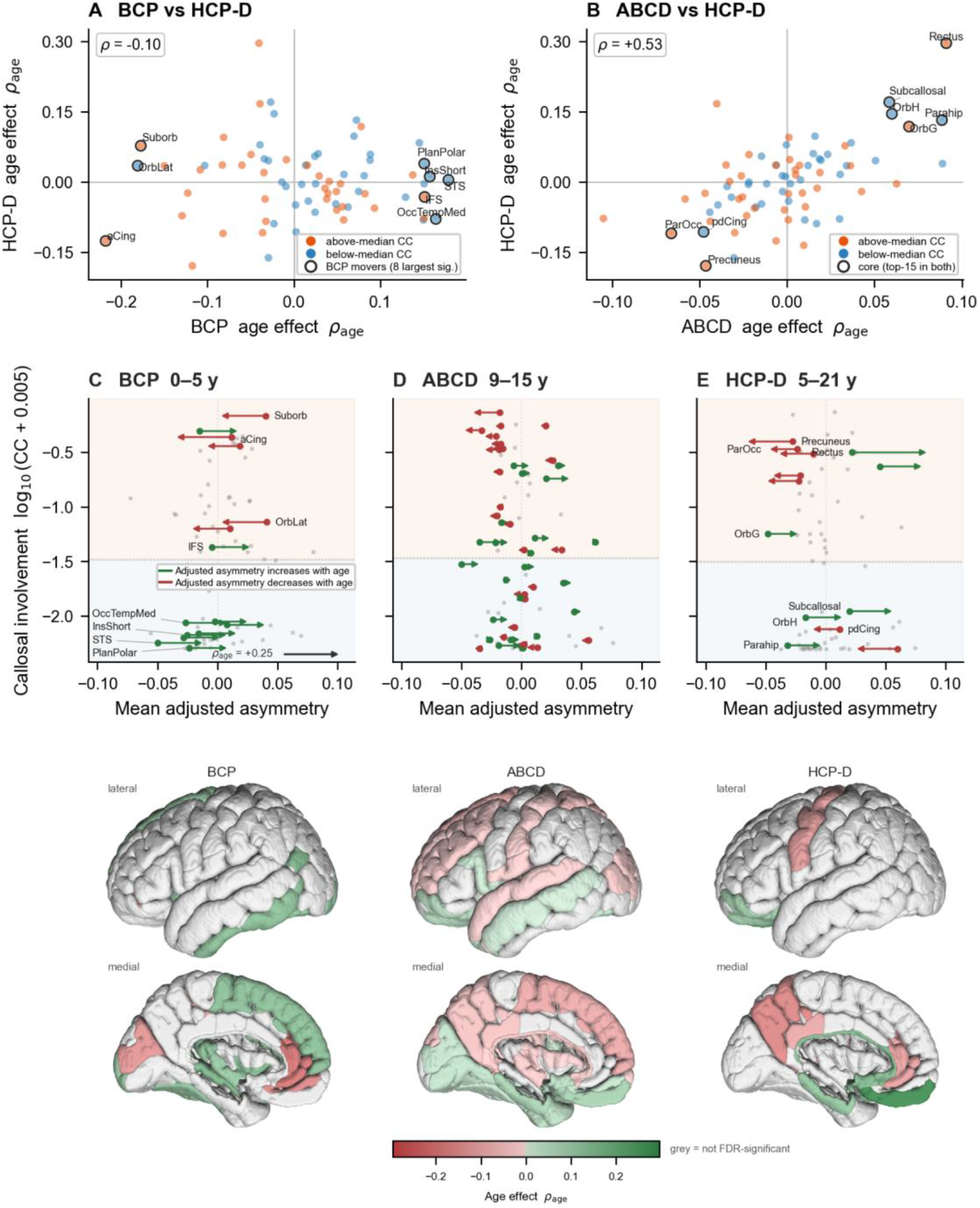
Regional age effects agree between the older cohorts and differ from BCP. Samples match Fig. 4; ρ_age_ denotes Spearman ρ(age, adjusted asymmetry). (A, B) Regional age effects in BCP and ABCD against HCP- D, colored by HCP-D’s median callosal split. Circles identify different sets: BCP’s eight largest significant age effects in A, and the eight regions shared by ABCD’s and HCP-D’s top-15 absolute age effects in B. (C–E) Regions positioned by mean adjusted asymmetry and callosal overlap. Arrows encode age effects on a common scale, not displacement in asymmetry units: green indicates increasing asymmetry magnitude, red decreasing magnitude. Arrows appear for false-discovery-rate (FDR)-significant regional age associations; gray dots indicate other regions. Background tints mark each cohort’s median callosal split. Labels identify selected regions in C and E. (Bottom) Left-hemisphere surface maps use the same diverging age-effect scale, with nonsignificant regions gray. Map correlations are descriptive and were not tested against a spatial null. Regional significance is calculated across sessions and may be anti-conservative in BCP and ABCD because participants contribute repeated scans. Reliability and membership checks are in the SI Appendix, section S7 and Table S8.

Eight regions were shared between ABCD’s and HCP-D’s 15 largest absolute age effects, including gyrus rectus, precuneus, subcallosal gyrus, and parahippocampal gyrus. This shared set was concentrated medially, with a median midline distance of 11.2 mm compared with 35.1 mm across all 74 regions. Seven of the eight remained when ABCD’s age effects were estimated from follow-up waves alone (SI Appendix, section S7).

Tract-level decomposition identifies the pathways contributing to these regional changes (Fig. 6). ABCD and HCP-D showed similar patterns across the displayed age-effect cells, including association-family raw absolute left–right differences and callosal overlap (ρ = +0.60), while BCP again showed little correspondence. This descriptive decomposition locates the change within specific pathways; it does not independently retest the regional interaction. The two older cohorts therefore share an anatomical pattern of change that differs from BCP’s, while the developmental and cohort contributions to that difference remain unresolved.

**Figure 6.**
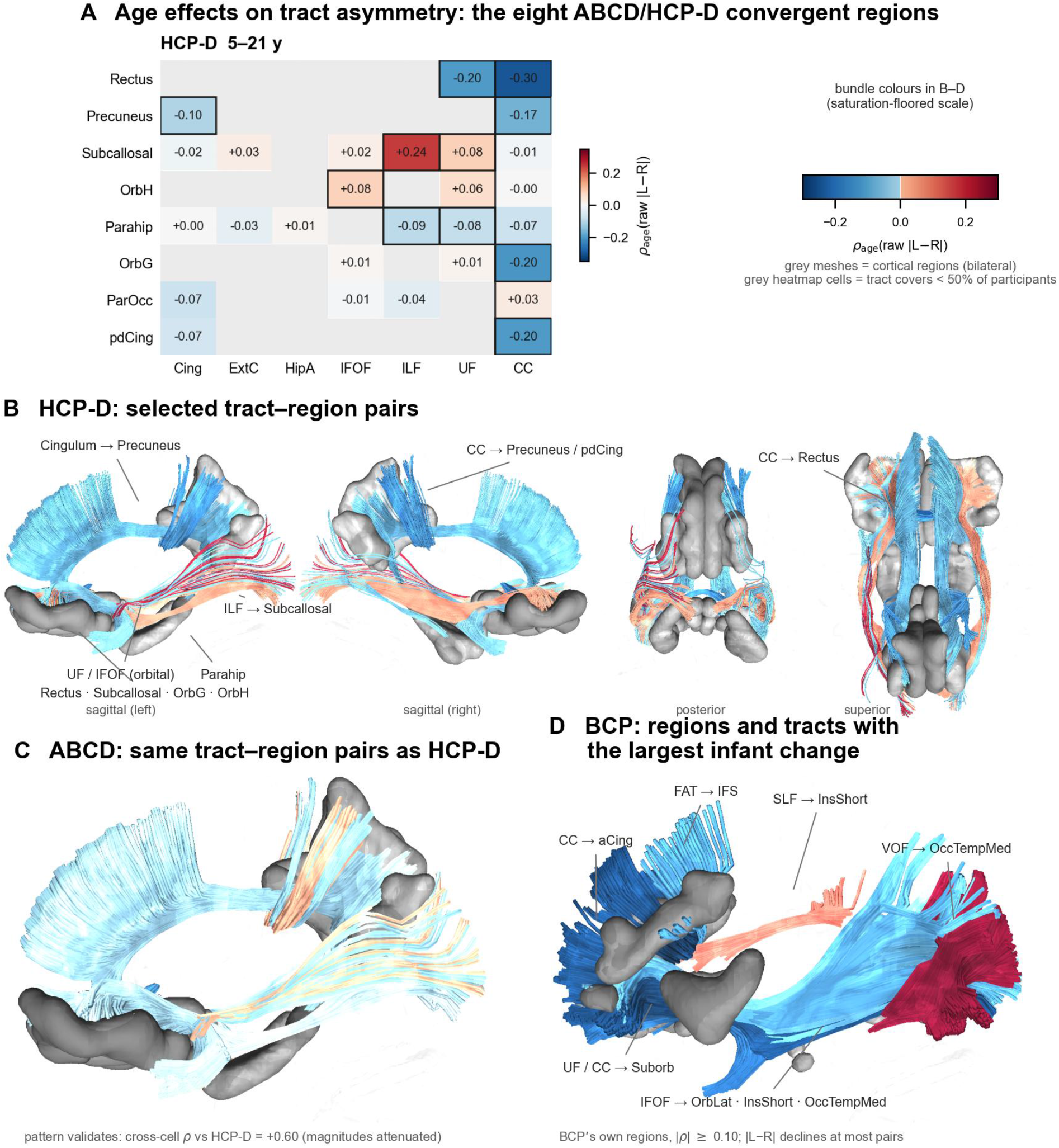
Tract contributions to regional changes in asymmetry. (A) HCP-D age correlations with raw absolute left–right overlap differences for the eight convergent regions in Fig. 5B. Association-family cells require coverage in at least 50% of participants; gray denotes lower coverage. The final column gives the age effect on callosal overlap itself. Outlines identify rendered cells. (B) Selected HCP-D tract–region pairs in four views. Gray meshes show cortical regions. Bundle colors use a saturation floor to keep weak effects visible; numerical magnitudes should be read from the heatmap. (C) The same tract–region pairs in ABCD. (D) Selected BCP pairs with absolute age correlations ≥0.10, drawn from its own largest changes. A callosal projection to lateral orbital cortex could not be rendered because template streamlines were absent even after four-voxel dilation. This decomposition uses raw asymmetry because the adjustment is defined only at region level. It describes anatomical contributions and does not independently test the regional interaction. Abbreviations and cross-cell comparisons are in the SI Appendix, Table S10 and section S7.

## Discussion

How asymmetric a cortical region is and how its asymmetry changes with age have different relationships with connectivity. More asymmetric regions had lower overall connectional density in all three cohorts, extending to cortical and non-cortical targets. In contrast to expectations from functional- lateralization studies (10), callosal involvement did not show a consistent inverse relationship with asymmetry magnitude. Instead, in both older cohorts, it distinguished the relationship between mean regional asymmetry and age-related change. The conditional regional relationship identified here would be missed by a static comparison alone (Figs. 3 and 4).

The distinction depends on what is measured. Functional studies have linked lateralization to preferential communication within a hemisphere and to larger-scale cortical organization (9, 11). Karolis and colleagues found that stronger functional lateralization accompanied lower probability of callosal connectivity (10). Our analysis measures left–right differences in association-tract overlap and the extent of callosal overlap with a region. These quantities do not directly measure functional dominance, axon number, or the probability used in that earlier study. The different findings therefore identify a need to connect structural and functional measures within the same developmental design. They do not overturn the functional observations. Likewise, the absence of a consistent static correlation does not establish that asymmetry and callosal involvement are independent.

Wu and colleagues linked callosal diffusion microstructure to subsequent left–right differences in cortical thickness and surface area and in subcortical volume (14). Here, regional asymmetry summarizes association tract connectivity by combining the magnitudes of tract-family differences, including pathways with opposite hemispheric directions (Fig. 1). In the older cohorts, greater mean adjusted asymmetry accompanied more positive age effects where callosal overlap was appreciable; this relationship was weak or reversed where overlap was low (Fig. 4D–E). Callosal involvement thus distinguished contrasting developmental patterns of tract connectivity. Their Destrieux analysis also linked age to thickness asymmetry in gyrus rectus and surface-area asymmetry in subcallosal gyrus (14), providing anatomical overlap with our eight-region set (Figs. 5 and 6).

The tract decomposition also clarifies what regional structural asymmetry means. Opposite hemispheric differences can coexist within one region, as illustrated by the arcuate and middle longitudinal fasciculi in planum temporale. Treating the region as uniformly leftward or rightward would lose that information. Measuring the magnitude of each family’s difference before averaging preserves this internal diversity. The resulting regional pattern is reproducible across cohorts, while the developmental pattern differs more substantially. Thus, a relatively stable arrangement of more and less asymmetric regions can coexist with regional differences in ongoing change.

Existing accounts of specialization provide context but address different scales of explanation. Conduction-delay arguments concern the costs of interhemispheric communication, especially as brains enlarge across species (5, 6, 8); human allometric findings also relate brain size to connectional organization (7). A regional association within the human cortex is not a direct test of those evolutionary accounts. Similarly, structural overlap cannot distinguish the integrative and inhibitory functions attributed to callosal pathways (12, 13). Our findings instead identify a developmental relationship that these functional and evolutionary accounts could help explain.

The developmental evidence has two components. In BCP, regional asymmetry converged toward the mean, narrowing its between-region spread (Fig. 4A,C). HCP-D, which spans a broader developmental age range, provides the primary evidence that callosal involvement conditions the relationship between mean asymmetry and age effects (Fig. 4B,E). ABCD provides directionally consistent support within its narrower age window (Fig. 4B,D). Early cortical and white-matter asymmetries are well established (2–4); the present results concern their regional organization and subsequent change, not their first appearance. Agreement at matched ages (Fig. 4G) strengthens the comparison between the two older cohorts. The BCP contrast, however, remains confounded with acquisition differences and age-dependent measurement properties. Neither the cohort ordering nor the within-cohort analyses establish a developmental onset or progressive strengthening of the interaction.

In the older cohorts, greater average asymmetry went with a stronger tendency toward increasing asymmetry with age where callosal overlap was appreciable; this relationship was weak or reversed where overlap was low (Fig. 4D–E). Callosal involvement therefore helps explain how the regional pattern of asymmetry develops, even though it shows no consistent relationship with asymmetry magnitude alone (Fig. 3). In BCP, regional asymmetry instead converged without detected callosal moderation (Fig. 4A–C). Biologically, the contrast raises the possibility that broad convergence early in life gives way to regional asymmetry development associated with interhemispheric connectivity. The cohorts show these contrasting patterns, but do not identify when the relationship with callosal involvement changes.

Causal explanations remain open. Experiments in mouse somatosensory cortex show that neuronal activity contributes to the development and maintenance of callosal projections (18), making activity-dependent refinement one plausible mechanism to investigate. Alternatively, regional developmental programs could influence both callosal connectivity and asymmetry trajectories. Human genetic associations with structural or functional asymmetry support considering developmental molecular influences (19, 20), without establishing shared regulation of the two measures studied here. A regional index of the age association with mean bilateral association overlap did not account for the interaction (SI Appendix, section S9), but it captures only one aspect of maturation. Neither this control nor the observational design distinguishes the causal accounts or establishes temporal precedence.

The distinction may also help frame studies of neurodevelopmental conditions. Large-scale analyses have identified regional alterations of structural asymmetry in autism spectrum disorder (21), schizophrenia (22), attention-deficit/hyperactivity disorder (23) and obsessive-compulsive disorder (24), and altered structural and functional asymmetry has been reported in genetic neurodevelopmental syndromes such as Williams syndrome (20, 25). Such differences need not be accompanied by a simple reduction or increase in callosal involvement at one age. A useful next question is whether the relationship between connectivity and asymmetry trajectories differs between groups. Answering it requires longitudinal clinical data and measures of function; the present study does not establish a disease marker or a basis for intervention.

Several limits determine how the developmental findings should be used. ABCD’s longitudinal interaction agreed in magnitude with the cross-sectional estimate but did not exceed the spatial null, and its inference depended on the variance estimator. Individual change was dominated by measurement error under the shared-wave model, limiting interpretation to region means. Change maps were less reliable than mean-asymmetry maps, especially in BCP and HCP-D. Tractography also carries ambiguity in pathway reconstruction (26): use of two tracking procedures avoids direct mathematical coupling between asymmetry and degree but does not eliminate shared imaging biases.

Midline position is similarly inseparable from aspects of callosal anatomy. Finally, the different ABCD transition estimates and BCP’s disagreement with the older cohorts remain unexplained. Whether the contrast between BCP and the older cohorts reflects a transition after which the regional organization of asymmetry and its callosal relationship begin to take their later form is a question for denser longitudinal sampling across childhood.

The main implication of the results reported here is that the connectivity of an asymmetric region cannot be summarized by a single static relationship. Lower overall connectional density accompanies greater structural asymmetry, while callosal involvement distinguishes its developmental pattern in the older cohorts. Studying magnitude and change together provides a more specific account of how hemispheric differences are organized within the developing connectome, and a basis for asking how alterations of these developmental relationships may contribute to neurodevelopmental and psychiatric conditions.

## Materials and methods

### Cohorts and analysis units

We analyzed BCP (450 sessions, 429 with usable age; 0–5.4 years) (27), ABCD (19,569 sessions from 10,652 participants across three waves; 8.9–15.7 years) (28), and HCP-D (636 participants; 5.6–21.9 years) (29, 30). Analyses were conducted separately within each cohort across 74 homotopic cortical region pairs; no brain-derived values were pooled across cohorts. BCP and ABCD include repeated sessions, while the HCP-D sample analyzed here is cross-sectional. Sample definitions and extended acquisition and processing details are in the SI Appendix, section S1 and Table S1. All data were obtained in de-identified form from the NIMH Data Archive under approved data use certifications; the original studies obtained informed consent from participants or their guardians under their respective institutional review boards. The present study is a secondary analysis of these de-identified, publicly available data and involved no new data collection or interaction with human participants. It was conducted under the University of Pittsburgh Institutional Review Board protocol STUDY19040351 (approved June 28, 2019), which covers image analysis of publicly available imaging data.

Connectional-density analyses used 636 HCP-D participants, 446 BCP sessions with usable asymmetry, and a separate subsample of 1,000 ABCD baseline participants selected without replacement, one per family, with allocation proportional to site (SI Appendix, section S1).

For ABCD, ComBat (31) was applied separately to two session-by-region matrices: raw absolute asymmetry and total association overlap across the 74 region pairs. Batch was defined by the acquisition site recorded for each session (22 sites). Continuous age in years and categorical sex were included as biological covariates to preserve their modeled effects. The three waves were pooled without a separate wave covariate; repeated sessions entered as separate rows, without participant-specific effects or longitudinal harmonization. Missing regional values were temporarily imputed using regional medians and re-masked after harmonization; harmonized values were constrained to be nonnegative. The within-session overlap adjustment was then recomputed from the harmonized measures. Callosal overlap was not harmonized in ABCD. BCP association and callosal overlap were harmonized with ComBat across scanner type, preserving age and sex and adjusting location only (SI Appendix, section S1).

### Connectivity and asymmetry measures

Tractography used DSI Studio and preprocessed Fiber Data Hub data (32), reconstructed with generalized q-sampling imaging (33). Atlas-based recognition reconstructed named association pathways (long-range intrahemispheric pathways, as distinct from commissural and projection pathways, grouped into 12 bilateral tract families; SI Appendix, section S1) and callosal segments (34). For each tract and Destrieux cortical region (35), the intersect ratio was the fraction of regional voxels whose tract density exceeded 0.5% of that tract’s peak. A separate coverage rule excluded family– region cells when the sum of left and right intersect ratios was ≤0.005. These are distinct voxel-level and cell-level thresholds (SI Appendix, section S1).

For each covered tract family, we calculated the absolute difference between its left and right regional overlap. We averaged these absolute differences over covered families, so opposing directions could not cancel and regions with more families did not receive greater weight simply through summation. Raw asymmetry depends strongly on total association overlap. Within each session, we therefore regressed the family-mean absolute difference across regions on log total association overlap and retained the residual. This total-overlap-adjusted asymmetry, abbreviated adjusted asymmetry, retains the units of the absolute difference; it is not divided by the sum of left and right overlap. Regional means were then calculated across sessions. Positive residuals indicate greater asymmetry than predicted by total overlap, not statistical significance or a particular hemispheric direction. Alternative adjustment and coverage analyses are in the SI Appendix, sections S2 and S3.

Callosal involvement was the summed overlap of callosal segments with each region pair, left unadjusted. Overall connectional density was obtained in every cohort from separate whole-brain tractography and a symmetric region-to-region matrix built with one pipeline (SI Appendix, section S1). An edge’s weight was the number of streamlines passing through both regions, with no connectivity threshold, so an edge is zero only when no reconstructed streamline links the two regions. Cortical degree was the mean edge weight to 147 other cortical targets, averaged over the left and right copies of each region. Non-cortical degree averaged over 18 bilateral subcortical and cerebellar gray-matter targets. Node definitions, tracking parameters, matched-target comparisons, and sensitivity analyses are in the SI Appendix, sections S1 and S4.

### Developmental estimates and models

For each region, the cross-sectional age effect (ρ_age_) was the Spearman correlation between age and adjusted asymmetry across sessions within cohort. It measures the direction and strength of an age association. Mean adjusted asymmetry is each region’s cohort average across sessions within the sampled age window. We correlated regional mean asymmetry with age effects and assessed dependence arising from shared data by estimating the two quantities in disjoint subject halves over 200 replications. We also compared between-region dispersion in age bins matched on session count, with correction for sampling error.

The primary regional model included mean asymmetry, callosal involvement, and their interaction as predictors of the age effect, with standardized coefficients. Callosal involvement entered the model without transformation. Mean adjusted asymmetry and callosal overlap were z-scored across the 74 regions before multiplication to form the interaction term; the product was not standardized again.

Logarithmic callosal scales were used only for visualization. A positive interaction means that greater callosal involvement shifts the mean-asymmetry–age-effect relationship in a positive direction. Median splits illustrate the model but do not define the inferential test. We also fitted sensitivity models for curvature, influence, midline distance, and tract coverage, and repeated the HCP-D model within ABCD’s age range (SI Appendix, sections S2 and S5).

For ABCD, longitudinal change was the difference in adjusted asymmetry between consecutive visits, averaged across visit pairs. The cohort-mean regional asymmetry served as the predictor, avoiding direct coupling to each individual’s earlier measurement. Consecutive two-year transitions were primary; pooling all pairs, including four-year intervals, was a sensitivity analysis. Transition comparisons, window-width analyses, and the shared-wave measurement-error model are described in the SI Appendix, section S6.

### Inference and anatomical comparisons

We used Spearman correlations for bivariate regional and within-participant relationships. Regional spatial checks used 10,000 spin rotations preserving spatial autocorrelation (36), with empirical exceedance P values. For the interaction models, the regional outcome map is rotated while mean asymmetry, callosal involvement, and their product remain fixed; the null therefore breaks the outcome’s spatial relationship with all model predictors. It therefore provides a spatial-specificity check rather than a standalone test of moderation. Ordinary, precision-weighted, HC3-robust, and bootstrap estimates were distinguished where reported; the latter procedures do not themselves account for spatial dependence. Detailed specifications are in the SI Appendix, section S2.

Tract-direction consistency was assessed across participants within covered family–region cells, with multiplicity correction. Map reliability used subject split halves and Spearman–Brown correction; cross- map correlations were additionally corrected for both maps’ reliability. The eight convergent regions were the intersection of ABCD’s and HCP-D’s top-15 absolute age effects. Tract-level attribution used raw absolute differences because the regional adjustment has no family-level counterpart. The map comparisons and tract attribution are descriptive. Session-based regional significance in BCP and ABCD can be anti-conservative because participants contribute repeated scans; Fig. 5 is interpreted accordingly. Details and sex comparisons are in the SI Appendix, sections S3, S7 and S8.

### Data, Materials, and Software Availability

BCP data are available from the NIMH Data Archive (NDA), collection 2848 (datasets 24282, 33498 and 29633). HCP-D data are available from NDA collection 2846 (datasets 35393, 35406 and 37361). ABCD data (release 5.1) are available from NDA (https://doi.org/10.15154/z563-zd24) under a data use certification. Preprocessed fiber data were obtained from the Fiber Data Hub (https://brain.labsolver.org), and DSI Studio (build “Hou,” 4 August 2024) is freely available at https://dsi-studio.labsolver.org. Analysis code and the per-region summary values underlying all figures and SI tables (for each of the 74 homotopic region pairs in each cohort: mean raw and adjusted asymmetry, total association overlap, callosal overlap, cortical and non-cortical degree, cross-sectional age effects and ABCD region-mean change) are deposited at Zenodo (https://doi.org/10.5281/zenodo.23736810). The deposit is embargoed until publication of this article.

Software versions are listed in SI Appendix, section S1.

## Author contributions

Siva Venkadesh: Conceptualization, Investigation, Methodology, Formal analysis, Resources, Writing – original draft. Fang-Cheng Yeh: Methodology, Resources, Funding acquisition, Writing – review and editing. Patrick R. Hof: Conceptualization, Investigation, Methodology, Writing – review and editing.

Julie R. Korenberg: Conceptualization, Investigation, Methodology, Writing – review and editing.

## Funding

This work was supported by NIH R01NS120954 (F.-C.Y.) and by a subaward to F.-C.Y. from NIH R01MH134004 (ENIGMA consortium, University of Southern California), which partly supported S.V.

## Acknowledgments

Computational resources were provided by the Pittsburgh Supercomputing Center through NSF ACCESS allocations MED240021, MED230052 and CIS200026.

We thank the participants, families and study teams of the UNC/UMN Baby Connectome Project (BCP), the Adolescent Brain Cognitive Development (ABCD) Study® and the Human Connectome Project in Development (HCP-D) for making these data available. BCP data collection was supported by the National Institute of Mental Health under award U01MH110274, and HCP-D data collection under award U01MH109589. BCP and HCP-D data were obtained from the NIMH Data Archive (NDA; collections 2848 and 2846). This manuscript reflects the views of the authors and may not reflect the opinions or views of the NIH or of the investigators who submitted the original data to NDA.

Data used in the preparation of this article were obtained from the Adolescent Brain Cognitive Development (ABCD) Study (https://abcdstudy.org), held in the NIMH Data Archive (NDA). This is a multisite, longitudinal study designed to recruit more than 10,000 children aged 9–10 and follow them over 10 years into early adulthood. The ABCD Study® is supported by the National Institutes of Health and additional federal partners under award numbers U01DA041048, U01DA050989, U01DA051016, U01DA041022, U01DA051018, U01DA051037, U01DA050987, U01DA041174, U01DA041106, U01DA041117, U01DA041028, U01DA041134, U01DA050988, U01DA051039, U01DA041156, U01DA041025, U01DA041120, U01DA051038, U01DA041148, U01DA041093, U01DA041089, U24DA041123 and U24DA041147. A full list of supporters is available at https://abcdstudy.org/about/federal-partners/. A listing of participating sites and a complete listing of the study investigators can be found at https://abcdstudy.org/consortium_members/. ABCD consortium investigators designed and implemented the study and/or provided data but did not necessarily participate in the analysis or writing of this report. This manuscript reflects the views of the authors and may not reflect the opinions or views of the NIH or ABCD consortium investigators. The ABCD data repository grows and changes over time. The ABCD data used in this report came from NDA Study 2313, Annual Release 5.1 (https://doi.org/10.15154/z563-zd24).

AI tools were used to assist with writing, development of analysis and visualization scripts, and technical critique. The authors made all scientific judgments, verified all AI-assisted outputs and take full responsibility for the final manuscript.

## Competing interests

The authors declare no competing interests.

## Notes

https://doi.org/10.5281/zenodo.23736810

### Competing Interest Statement

The authors have declared no competing interest.

## References

1. O. Güntürkün, F. Ströckens, S. Ocklenburg, Brain lateralization: A comparative perspective. Physiological Reviews 100, 1019–1063 (2020). 10.1152/physrev.00006.2019

2. G. Li et al., Mapping longitudinal hemispheric structural asymmetries of the human cerebral cortex from birth to 2 years of age. Cerebral Cortex 24, 1289–1300 (2014). 10.1093/cercor/bhs413

3. L. Z. J. Williams et al., Structural and functional asymmetry of the neonatal cerebral cortex. Nature Human Behaviour 7, 942–955 (2023). 10.1038/s41562-023-01542-8

4. J. Dubois et al., Structural asymmetries in the infant language and sensori-motor networks. Cerebral Cortex 19, 414–423 (2009). 10.1093/cercor/bhn097

5. J. L. Ringo, R. W. Doty, S. Demeter, P. Y. Simard, Time is of the essence: A conjecture that hemispheric specialization arises from interhemispheric conduction delay. Cerebral Cortex 4, 331–343 (1994). 10.1093/cercor/4.4.331

6. J. K. Rilling, T. R. Insel, Differential expansion of neural projection systems in primate brain evolution. NeuroReport 10, 1453–1459 (1999). 10.1097/00001756-199905140-00012

7. J. Hänggi, L. Fövenyi, F. Liem, M. Meyer, L. Jäncke, The hypothesis of neuronal interconnectivity as a function of brain size—a general organization principle of the human connectome. Frontiers in Human Neuroscience 8, 915 (2014). 10.3389/fnhum.2014.00915

8. D. J. Ardesch et al., Scaling principles of white matter connectivity in the human and nonhuman primate brain. Cerebral Cortex 32, 2831–2842 (2022). 10.1093/cercor/bhab384

9. S. J. Gotts et al., Two distinct forms of functional lateralization in the human brain. Proceedings of the National Academy of Sciences USA 110, E3435–E3444 (2013). 10.1073/pnas.1302581110

10. V. R. Karolis, M. Corbetta, M. Thiebaut de Schotten, The architecture of functional lateralisation and its relationship to callosal connectivity in the human brain. Nature Communications 10, 1417 (2019). 10.1038/s41467-019-09344-1

11. L. Labache, T. Ge, B. T. T. Yeo, A. J. Holmes, Language network lateralization is reflected throughout the macroscale functional organization of cortex. Nature Communications 14, 3405 (2023). 10.1038/s41467-023-39131-y

12. J. S. Bloom, G. W. Hynd, The role of the corpus callosum in interhemispheric transfer of information: Excitation or inhibition? Neuropsychology Review 15, 59–71 (2005). 10.1007/s11065-005-6252-y

13. L. J. van der Knaap, I. J. M. van der Ham, How does the corpus callosum mediate interhemispheric transfer? A review. Behavioural Brain Research 223, 211–221 (2011). 10.1016/j.bbr.2011.04.018

14. X. Wu et al., Developing brain asymmetry shapes cognitive and psychiatric outcomes in adolescence. Nature Communications 16, 4480 (2025). 10.1038/s41467-025-59110-9

15. S. Ocklenburg, P. Friedrich, O. Güntürkün, E. Genç, Intrahemispheric white matter asymmetries: The missing link between brain structure and functional lateralization? Reviews in the Neurosciences 27, 465–480 (2016). 10.1515/revneuro-2015-0052

16. M. Thiebaut de Schotten, et al., A lateralized brain network for visuospatial attention. Nature Neuroscience 14, 1245–1246 (2011). 10.1038/nn.2905

17. M. Catani et al., Symmetries in human brain language pathways correlate with verbal recall. Proceedings of the National Academy of Sciences USA 104, 17163–17168 (2007). 10.1073/pnas.0702116104

18. C.-L. Wang et al., Activity-dependent development of callosal projections in the somatosensory cortex. Journal of Neuroscience 27, 11334–11342 (2007). 10.1523/JNEUROSCI.3380-07.2007

19. Z. Sha et al., The genetic architecture of structural left–right asymmetry of the human brain. Nature Human Behaviour 5, 1226–1239 (2021). 10.1038/s41562-021-01069-w

20. D. L. Mills, L. Dai, J. R. Korenberg, Brain lateralization for perceiving direction of motion is reversed in Williams syndrome and related to BUD23. Scientific Reports 15, 19772 (2025). 10.1038/s41598-025-94742-3

21. M. C. Postema et al., Altered structural brain asymmetry in autism spectrum disorder in a study of 54 datasets. Nature Communications 10, 4958 (2019). 10.1038/s41467-019-13005-8

22. D. Schijven et al., Large-scale analysis of structural brain asymmetries in schizophrenia via the ENIGMA consortium. Proceedings of the National Academy of Sciences USA 120, e2213880120 (2023). 10.1073/pnas.2213880120

23. M. C. Postema et al., Analysis of structural brain asymmetries in attention-deficit/hyperactivity disorder in 39 datasets. Journal of Child Psychology and Psychiatry 62, 1202–1219 (2021). 10.1111/jcpp.13396

24. X.-Z. Kong et al., Mapping cortical and subcortical asymmetry in obsessive-compulsive disorder: Findings from the ENIGMA Consortium. Biological Psychiatry 87, 1022–1034 (2020). 10.1016/j.biopsych.2019.04.022

25. M. A. Eckert et al., Anomalous sylvian fissure morphology in Williams syndrome. NeuroImage 33, 39–45 (2006). 10.1016/j.neuroimage.2006.05.062

26. K. H. Maier-Hein et al., The challenge of mapping the human connectome based on diffusion tractography. Nature Communications 8, 1349 (2017). 10.1038/s41467-017-01285-x

27. B. R. Howell et al., The UNC/UMN Baby Connectome Project (BCP): An overview of the study design and protocol development. NeuroImage 185, 891–905 (2019). 10.1016/j.neuroimage.2018.03.049

28. B. J. Casey et al., The Adolescent Brain Cognitive Development (ABCD) study: Imaging acquisition across 21 sites. Developmental Cognitive Neuroscience 32, 43–54 (2018). 10.1016/j.dcn.2018.03.001

29. L. H. Somerville et al., The Lifespan Human Connectome Project in Development: A large-scale study of brain connectivity development in 5–21 year olds. NeuroImage 183, 456–468 (2018). 10.1016/j.neuroimage.2018.08.050

30. M. P. Harms et al., Extending the Human Connectome Project across ages: Imaging protocols for the Lifespan Development and Aging projects. NeuroImage 183, 972–984 (2018). 10.1016/j.neuroimage.2018.09.060

31. J.-P. Fortin et al., Harmonization of multi-site diffusion tensor imaging data. NeuroImage 161, 149–170 (2017). 10.1016/j.neuroimage.2017.08.047

32. F.-C. Yeh, DSI Studio: An integrated tractography platform and fiber data hub for accelerating brain research. Nature Methods 22, 1617–1619 (2025). 10.1038/s41592-025-02762-8

33. F.-C. Yeh, V. J. Wedeen, W.-Y. I. Tseng, Generalized q-sampling imaging. IEEE Transactions on Medical Imaging 29, 1626–1635 (2010). 10.1109/TMI.2010.2045126

34. F.-C. Yeh, Population-based tract-to-region connectome of the human brain and its hierarchical topology. Nature Communications 13, 4933 (2022). 10.1038/s41467-022-32595-4

35. C. Destrieux, B. Fischl, A. Dale, E. Halgren, Automatic parcellation of human cortical gyri and sulci using standard anatomical nomenclature. NeuroImage 53, 1–15 (2010). 10.1016/j.neuroimage.2010.06.010

36. A. F. Alexander-Bloch et al., On testing for spatial correspondence between maps of human brain structure and function. NeuroImage 178, 540–551 (2018). 10.1016/j.neuroimage.2018.05.070

