## Supplementary material for "Cortical asymmetry and its developmental change have distinct relationships with brain connectivity": SI Appendix

This PDF file includes: Supplementary text (sections S1 to S10), Figs. S1 to S8, Tables S1 to S10, and SI references

This SI Appendix accompanies “Cortical asymmetry and its developmental change have distinct relationships with brain connectivity.” It contains extended methods, supporting results, and numerical summaries. References in this appendix are numbered independently of the main manuscript.

### S1 Cohorts and connectome construction

#### Samples and preprocessing

The three cohorts were analyzed separately. BCP (1) samples infancy and early childhood; ABCD (2) provides repeated measurements across three waves; and HCP-D (3, 4) spans the widest age window. HCP-D encompasses ABCD's age range, so the cohorts should not be treated as three ordered developmental stages. All reported regional relationships use the 74 homotopic cortical pairs within a cohort, without pooling brain-derived values across cohorts.

Table S1 distinguishes the level sample, which requires usable asymmetry and callosal measurements but no age record, from the age-effect sample. BCP contributes 450 sessions from 240 participants to level analyses and 429 sessions from 232 participants to age analyses. Its sex-stratified sample includes 435 sessions, 213 male and 222 female. Per-region sample sizes vary with tract coverage: 207–450 in BCP, 14,070–19,569 in ABCD, and 278–636 in HCP-D. The joint ABCD sample comprises 19,569 sessions from 10,652 participants. Sessions, participants, and visit pairs are distinct counting units throughout.

**Table S1 Cohort samples used in the analyses**

| Quantity | BCP | ABCD | HCP-D |
|---|---|---|---|
| Observed age range in years | 0–5.4 | 8.9–15.7 | 5.6–21.9 |
| Sessions for level analyses | 450 | 19,569 | 636 |
| Sessions with usable age | 429 | 19,569 | 636 |
| Unique participants | 240 (level); 232 (age) | 10,652 | 636 |
| Repeated measurements in this analysis | Yes | Three waves | No |
| Sessions with a sex record | 435 | 19,569 | 636 |
| Sex sample | 213 male; 222 female | 10,237 male; 9,332 female | 292 male; 344 female |
| Per-region sample range by coverage | 207–450 | 14,070–19,569 | 278–636 |
| Homotopic cortical pairs | 74 | 74 | 74 |

Counts refer to the analysis samples, not the full cohort releases. The ABCD asymmetry-only sample has 19,605 sessions from 10,662 participants; intersecting it with usable callosal overlap gives the 19,569-session, 10,652-participant sample reported here. BCP has 240 unique participants in the level sample and 232 in the age sample. Region-to-region matrices were generated for 636 HCP-D participants, 531 BCP sessions (446 with usable asymmetry), and 1,000 ABCD baseline participants; these are distinct from the full level and age-effect samples above.

ABCD data were harmonized across 22 acquisition sites with ComBat (5), separately for raw absolute asymmetry and the association-overlap measure across 74 regions, preserving continuous age and categorical sex. Waves were pooled without a wave covariate, and repeated sessions were separate rows without participant-specific effects. Regional median imputation was temporary; missing cells were re-masked, harmonized values were constrained to be nonnegative, and the within-session adjustment was recomputed. In ABCD, callosal overlap was not harmonized. BCP data were also harmonized with ComBat, across scanner type (two levels, with the Prisma fit scanner as reference), preserving continuous age at scan and categorical sex and adjusting location only. Harmonization was applied to each association and callosal tract separately, with regional values as features; unlike ABCD, BCP callosal overlap was therefore harmonized. Sessions missing scanner type, age or sex, regions with zero variance, and tracts with fewer than 10 usable sessions or a single scanner type were left unharmonized, and harmonized values were clipped to [0, 1]. Tractography used preprocessed fiber data from the Fiber Data Hub (6) and generalized q-sampling imaging reconstruction (7).

BCP data were obtained from the NIMH Data Archive (NDA), collection 2848, package 1225906 (datasets 24282, 33498, and 29633; submissions 23233, 31933, and 28093). HCP-D used NDA collection 2846, datasets 35393, 35406, and 37361. ABCD used release 5.1. DSI Studio “Hou,” build 4 August 2024, was used for named-tract and whole-brain tracking. Analysis used Python 3.8.8, NumPy 1.24.4, SciPy 1.10.1, pandas 1.3.3, statsmodels 0.14.1, Matplotlib 3.7.5, and nilearn 0.8.0. Reconstruction and tracking were in native space; BCP six-shell GQI used a sampling-length ratio of 1.25. Data were AC–PC aligned and registered to ICBM152 using DSI Studio, with a cached per-session warp. Level analyses required tract-to-region and callosal measurements and at least one association family above the bilateral 0.005 coverage threshold. BCP retained visits 1–9 and preferred the six-shell reconstruction over the 79-direction reconstruction when both existed. ABCD used one row per visit without participant effects.

#### Tract-to-region representation

Named pathways were reconstructed by atlas-based automatic tract recognition against a population-based tractography atlas (8). Association pathways were mapped to 12 bilateral tract families; In ABCD, subdivisions of a matched family are retained as separate families for the 74-region asymmetry measure and are combined by summation only for the tract-level decomposition (main Fig. 6 and Fig. S6). Corpus-callosum segments were analyzed separately. For each recognized tract, a tract-density image was compared with the aligned FreeSurfer Destrieux parcellation (9). The intersect ratio is the fraction of a region’s voxels in which tract density exceeds 0.5% of that tract’s peak density. Each tract–region value lies between zero and one.

The 148 cortical regions form 74 homotopic pairs. Each association-tract family has an explicit left–right homologue, permitting comparison of its regional overlap across hemispheres. The representation measures how much a named pathway overlaps a region; it does not identify all region-to-region edges. The association-overlap magnitude is the mean bilateral overlap (L + R) across covered association families. Callosal involvement is summed across corpus-callosum segments. Asymmetry is averaged across covered association families, as described in section S2.

The voxel threshold and the family-coverage threshold operate on different quantities. A voxel contributes to an intersect ratio when its density exceeds 0.5% of the tract's peak. A family–region cell is then masked as uncovered when its left and right intersect ratios sum to ≤0.005. The shared numerical value does not make these the same criterion.

#### Region-to-region representation

Whole-brain tractography was run in native subject space for every cohort with a single software build (DSI Studio "Hou," build 4 August 2024, also used for the project's tract-to-region data), separately from named-tract recognition, with the FreeSurferSeg parcellation (192 labels: 148 cortical, 18 subcortical and cerebellar gray-matter, and white-matter, ventricular and brain-stem labels that are not used as nodes) warped from template to subject space. One million streamlines were generated per session by deterministic tracking (anisotropy threshold 0.6 of the Otsu value; step size, turning angle, and anisotropy cut-off chosen automatically from each dataset; streamline length 30–200 mm). A matrix entry is the number of streamlines passing through both regions, stored once per unordered pair and mirrored; no connectivity threshold is applied, so an entry is zero only when no streamline links the two regions.

ABCD matrices were generated for 1,000 baseline participants drawn without replacement, one participant per family, with allocation proportional to site. BCP matrices cover 531 sessions, with duplicate acquisitions resolved to the six-shell run; 446 of these carry the asymmetry measure. HCP-D covers 636 participants. These streamline-count matrices are used for the primary connectional-density analyses.

The complete FreeSurferSeg matrix has 192 nodes: 148 cortical regions, 18 non-cortical gray-matter structures, and 26 excluded segments. The non-cortical targets are the thalamus, caudate, putamen, pallidum, hippocampus, amygdala, accumbens area, ventral diencephalon, and cerebellar cortex, bilaterally. This is the complete non-cortical target set used in this analysis. Excluded labels comprise white matter, ventricular and cerebrospinal-fluid spaces, vascular and unassigned segments, the optic chiasm, brain stem, and the two cortical "Unknown" labels. Brain stem is excluded as a mixed gray- and white-matter structure, not because it lacks gray matter.

#### Why the two representations are separated

Cortical degree requires region-to-region edges. An overlap-based surrogate would measure tract incidence and would share the same tract-by-region matrix used to construct asymmetry. The present comparison instead relates atlas-recognized tract asymmetry to a graph produced by whole-brain tracking (Fig. S1). This avoids direct mathematical coupling between the two measures, although biases shared through the diffusion data, reconstruction, or tractography can remain.

Under the primary pass-through definition, 2.4% (HCP-D), 0.6% (BCP), and 7.9% (ABCD) of cross-hemisphere session-by-region cells are zero, compared with 0.0% of within-hemisphere cells; under the endpoint definition the cross-hemisphere floor rises to 20%, 7%, and 25%. A zero records that no connecting streamline was reconstructed, rather than a continuous edge weight falling below an additional threshold.

**Figure S1 Two representations of structural connectivity**

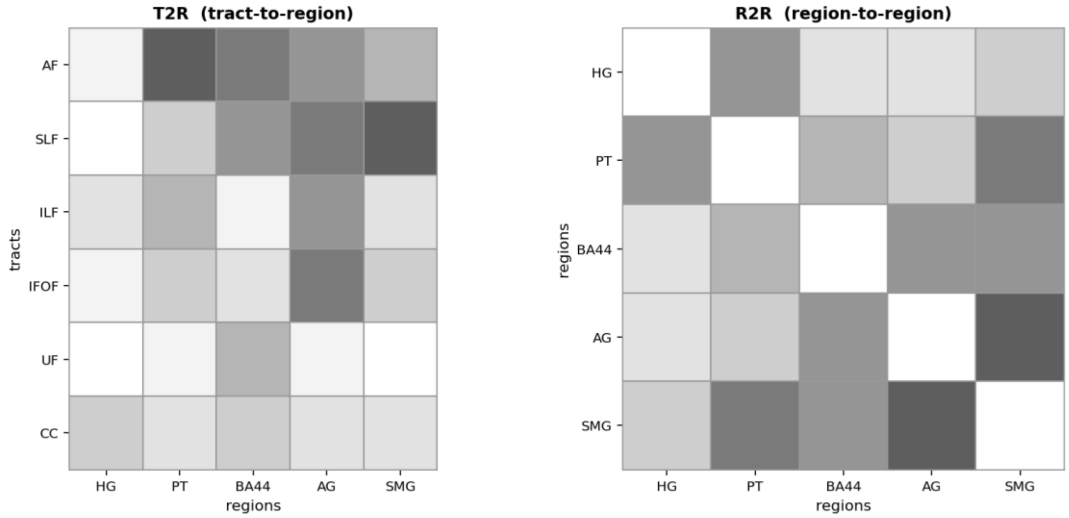


Illustrative tract-by-region and region-by-region matrices show the different units represented by the two constructions. Shading is schematic and does not encode observed data. The tract-by-region matrix records overlap of named pathways with cortical regions; the symmetric region-by-region matrix records connections between pairs of regions.

### S2 Measures and statistical analyses

#### Regional asymmetry and adjustment

For each covered family and region, asymmetry is the absolute left–right difference in intersect ratio. Absolute values are taken before averaging over families because opposing signed differences within a region would otherwise cancel. Averaging rather than summing avoids giving a region a larger value simply because it intersects more covered families.

Within each session, family-mean absolute difference is regressed across regions on log total association overlap. The residual is the total-overlap-adjusted asymmetry used in the main analyses. It is subsequently averaged over sessions to obtain regional means. This adjustment is denominator-free and retains the units of the absolute difference. Unlike a conventional lateralization ratio, it does not divide by the sum of left and right overlap. Negative residuals indicate less asymmetry than predicted from total overlap; they do not indicate rightward dominance or literal negative asymmetry. Raw and adjusted values are distinguished whenever both are shown.

Raw asymmetry correlates strongly with total overlap, motivating this adjustment (Table S2). Sensitivities replace the cohort-specific adjustment slope with a pooled slope while retaining the same fitting scheme, or additionally include the number of covered families. The former checks stability of regional ordering; the latter addresses residual dependence on tract coverage (section S3).

#### Degree and target sets

Cortical degree is the mean edge weight to 147 other cortical targets, including both hemispheres, averaged over a region's left and right copies. It is a weighted degree, rather than a thresholded edge count. However, because weights are nonzero only when a connection is reconstructed, the regional measure is related to the count of nonzero targets ($\rho$ = +0.76, +0.74, and +0.82 in HCP-D, BCP, and ABCD under the pass-through definition, and +0.90 to +0.95 under the endpoint definition, which yields a sparser graph). Mean and summed degree differ by a constant for a fixed target set and yield identical rank statistics.

Non-cortical degree averages over the 18 targets listed in section S1. These targets are excluded from cortical degree, so controlling non-cortical degree does not algebraically remove part of the outcome. The two degree measures nonetheless share substantial variance (section S4). A matched-count comparison recalculates cortical degree using 18 cortical targets; it addresses target number, while leaving anatomical differences between the two target sets.

#### Age effects and regional models

The cross-sectional age effect ($\rho_{age}$) is the Spearman correlation of age with adjusted asymmetry across sessions within a cohort. Mean adjusted asymmetry is each region's cohort average within the sampled age window. The across-region correlation between mean adjusted asymmetry and the age effect is denoted ρ in Fig. 4C–E and described explicitly in Table S5. Longitudinal change instead summarizes within-participant differences in adjusted asymmetry between visits (section S6).

Because both quantities are estimated from the same data, their association can be biased downward. The disjoint-subject control estimates mean asymmetry and age effects in separate subject halves over 200 replications. Between-region dispersion is compared in age bins matched on session count, and corrected for sampling error using each bin's own within-region variance.

The primary model regresses the regional age effect on mean asymmetry, callosal involvement, and their interaction across 74 regions. Mean adjusted asymmetry and untransformed cohort-mean callosal overlap are z-scored across the 74 regions before multiplication; their product enters without further standardization. Reported coefficients are standardized. Logarithmic callosal scales are used only for visualization. The main effect of mean asymmetry is evaluated at mean callosal involvement. Median and outer-tertile splits are used for illustration. The interaction coefficient, rather than a comparison of significance tests in the two groups, addresses the difference in slopes. Because callosal overlap is heavily skewed, plots label each group's mean overlap. The lines in main Fig. 4 are values from the interaction model, not separate regressions fitted within the displayed groups.

Sensitivity specifications add a mean-asymmetry quadratic and its interaction with callosal involvement, or a callosal quadratic. The influence trims summarized in Table S6 are applied to the linear primary model, removing the one to five regions with highest callosal overlap or regions identified by Cook's distance. Spatial-position checks adjust for distance from the midline or exclude medial-wall regions; for the exclusion, predictors are z-scored across all 74 regions and the model is fitted to the 57 retained regions. A matched-age analysis restricts HCP-D to ABCD's 8.9–15.7-year range. These checks assess different sources of sensitivity and are not interchangeable tests of the same assumption.

For the dispersion analysis, young and old age bins were matched on session count: BCP, 0.0–0.8 versus 1.5–5.4 years (144 sessions per bin, sampled with replacement from 144 and 149 available); ABCD, 8.9–10.3 versus 11.9–15.7 years (6,524 per bin); HCP-D, 5.6–12.2 versus 16.2–21.9 years (213 per bin). A session–region cell is undefined if no association family exceeds the bilateral 0.005 coverage threshold; regional means and correlations use available sessions. Cook's-distance exclusions use $>4/n$ from the linear primary model ($n = 74$); separate trims remove the one to five regions with highest callosal overlap. Midline distance is the mean absolute MNI x-coordinate of a region's Destrieux 2009 atlas voxels, averaged across homologues.

#### Spatial inference and other uncertainty estimates

Regional spatial relationships are assessed with 10,000 spin rotations (seed 0), reused across tests, following the spatial correspondence framework (10). Region centroids are unit-normalized mean vertex coordinates on the fsaverage5 left-hemisphere sphere using the Destrieux atlas distributed with nilearn 0.8.0. Each random three-dimensional rotation is converted to a one-to-one label permutation by minimum-total-angular-distance matching (Hungarian assignment). One hemisphere represents the

74 homotopic pairs; the medial wall is not a node. Bivariate tests permute the second map while holding the first fixed. Two-sided empirical P values are the fraction of rotations with an absolute statistic at least as large as observed, without a +1 correction. Zero exceedances are reported as $P < 0.001$ rather than $P = 0$. The endpoint-rule connectional-density sensitivity uses 2,000 rotations.

For interaction models, only the regional outcome map—age effect or region-mean within-participant change—is permuted; mean asymmetry, callosal overlap, and their product remain fixed. The null breaks the outcome's spatial relationship with all model predictors and is not an isolated moderation test conditional on retained main effects. Precision weights travel with the outcome in weighted spatial tests. Null standard deviations of the interaction coefficient are 0.1473 for HCP-D cross-sectional, 0.1924 for ABCD cross-sectional, and 0.1894 for ABCD within-participant change. These values refer to three analyses, rather than a comparison of the three cohorts.

Ordinary least-squares, heteroscedasticity-consistent HC3, precision-weighted, and bias-corrected bootstrap results are identified separately. Neither HC3 nor a bootstrap over regions accounts for spatial dependence. Subject splits address stability across participant samples; matched-age comparisons address replication across cohorts and acquisition settings. Adjustment for midline distance is descriptive because it can remove both genuine positional organization and distance-related measurement effects.

Main Fig. 1A uses kernel-density violins scaled independently to equal maximum width, not to sample size. Insets show the median, interquartile range, and the most extreme observations within 1.5 interquartile ranges of the nearer quartile. Values beyond the whiskers remain in the density but are not drawn individually. Main Fig. 2B shows HCP-D kernel-density violins with median bars and no extrema. In Fig. 2C, degree is expressed relative to each cohort's low-tertile median; lines join tertile medians and shaded bands show interquartile ranges across participants or BCP sessions. HCP-D additionally shows boxes and whiskers using the same quartile convention across 636 participants and jittered participant points; a small number of points above 1.4 are outside the displayed axis. Each cohort's low, middle, and high asymmetry tertiles contain 24–25 regions.

Main Fig. 4A uses bootstrap 95% percentile intervals: 250 session-resampling replicates for whole-cohort circles (ABCD capped at 6,000 sessions per replicate), 300 for BCP age-bin circles, 200 for ABCD wave circles, and 400 subject-pair replicates for within-subject squares. Disjoint-subject triangles instead show the median and 2.5th–97.5th percentiles across 200 random half-sample replicates; these summarize replicate spread rather than uncertainty of the full-sample estimate. In Fig. 4B, circles use region-level ordinary least-squares 95% t intervals (74 regions; 70 degrees of freedom), and within-subject squares use 300 subject-pair bootstrap replicates. Fig. 4F uses 2,000 subject-pair bootstrap replicates; Fig. 4G uses the same regional t-interval convention as the circles in B.

In main Fig. 4C–E, colored bands are model-implied 95% confidence bands (±1.96 SE) evaluated at each subgroup's mean callosal overlap using the fitted covariance matrix. They quantify uncertainty of the fitted line, not a prediction interval or the scatter of regional values. Black bivariate lines have no band. Insets use the same 95% convention for the fitted slope. Green/pink background shading in A and B marks the sign of the estimate, not uncertainty.

Precision weights are inverse squared standard errors of region-mean change: $w = 1/SEM^2$, where SEM is the standard deviation of individual changes divided by the square root of the number of covering wave-adjacent pairs. The standard errors underlying these weights correlate +0.505 with mean asymmetry, −0.026 with callosal overlap, and −0.011 with midline distance; the weights themselves correlate with the opposite signs. These pair-count standard errors do not account for participant clustering.

#### Sign consistency and reliability

For each family–region cell, the proportion of participants agreeing on the sign of the left–right difference is computed and assessed against chance, accounting for varying cell sample sizes. Cell results are aggregated over covered families where a region-level summary is used. For cells where fewer than approximately 80% of participants have bilateral coverage, sign consistency is also calculated only among those with both sides present. This distinguishes graded asymmetry from presence on one side and absence on the other.

Map reliability is estimated from subject split halves and corrected with the Spearman–Brown formula. Observed cross-map correlations are also disattenuated using the reliability of both maps. The adjustment applies to complete map comparisons rather than selectively to individual cells. Session-based regional age-association significance in BCP and ABCD does not fully account for repeated observations, so the arrows and selected-region displays are descriptive.

### S3 Tract direction and regional asymmetry

#### Tract-specific directions

At planum temporale in HCP-D, arcuate fasciculus overlap is greater on the left (median left 0.165, right 0.016; mean signed difference +0.13; n = 630), while middle longitudinal fasciculus overlap is greater on the right (0.058 and 0.150; mean signed difference −0.09; n = 633). The arcuate pattern is consistent with prior tractography describing leftward asymmetry of its direct segment (11). These are tract-specific observations, not evidence that every pathway within planum temporale has the same direction.

Across all 74 regions, the tract family with the highest sign consistency in each region has a median sign consistency of 0.911, and all regions exceed chance after Bonferroni correction across all 332 tested family–region cells. When signed family differences are instead averaged at region level, 38 of 74 regions are leftward, illustrating the loss of a common directional interpretation after aggregation.

BA44 provides another example of opposing directions in the same participants: arcuate fasciculus is leftward in 0.685 of covered participants and superior longitudinal fasciculus is rightward in 0.918. Prior work documents lateralized parietofrontal white-matter networks (12). Figure S2 summarizes 14 covered cells from a four-tract by five-region grid as descriptive examples of tract-specific directions.

**Figure S2 Tract-specific asymmetry directions**

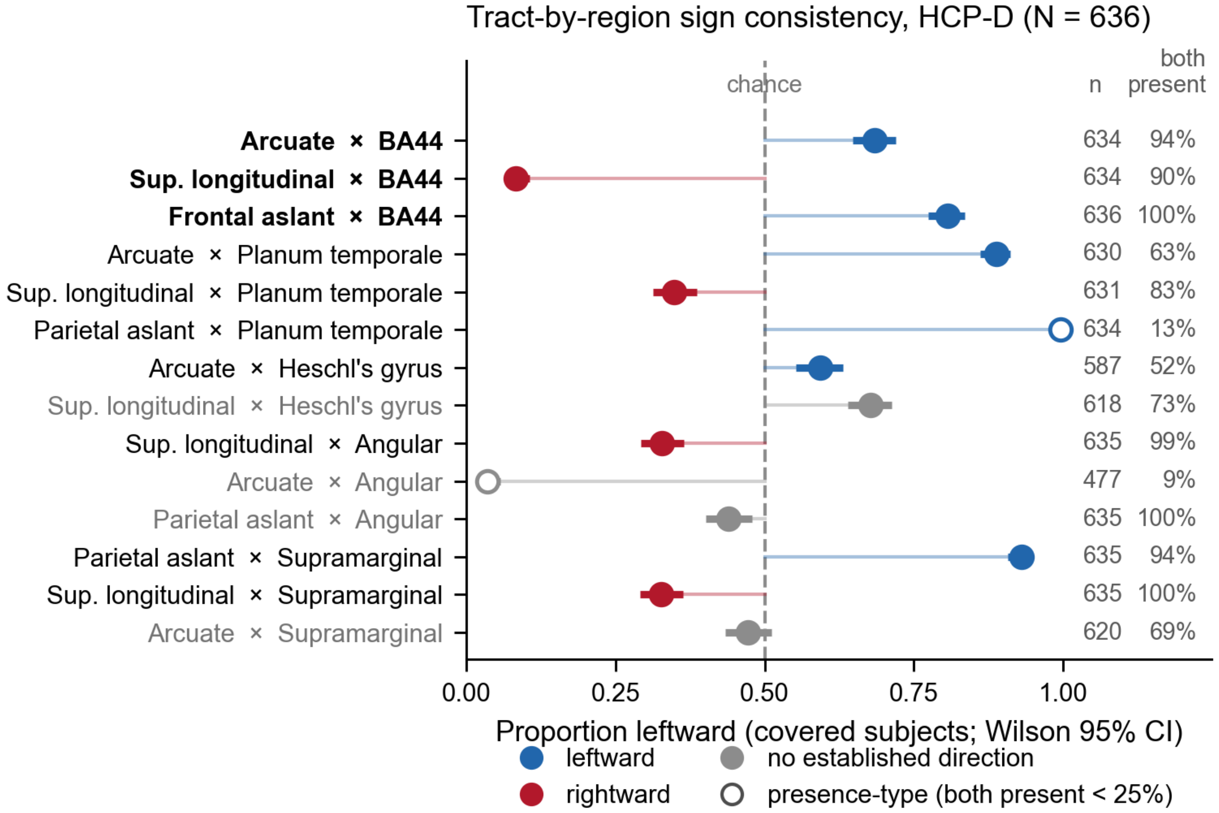


Proportion of covered participants with leftward asymmetry for the 14 covered tract–region cells in the four-tract by five-region grid. Markers show the observed proportion, with Wilson 95% confidence intervals; the chance value is 0.5. Blue and red highlight selected leftward and rightward patterns; gray denotes cells without an assigned reference direction. This display is descriptive and does not establish literature concordance for every cell. Covered sample sizes and the percentage with both sides present are printed for each cell. Open markers identify cells with bilateral presence below 25%, where the result largely reflects presence on one side rather than a graded difference between two present tracts. BA44 is included here but not in main Fig. 1B's magnitude-selected set: its regional adjusted-asymmetry ranks are 61, 47, and 33 of 74 in BCP, ABCD, and HCP-D, and its residual is not positive in any cohort.

### Regional ordering and adjustment sensitivities

The 29 positive-residual regions in HCP-D define the common display order in main Fig. 1C and Fig. S3B; Fig. S3A shows the 31 positive-residual regions in BCP in BCP order. Among the highest-ranking regions across cohorts are the anterior intermediate parietal sulcus, the vertical and horizontal rami of the anterior lateral fissure, transverse temporal sulcus, planum temporale, and orbital inferior frontal gyrus. The anterior intermediate parietal sulcus lies between the supramarginal and angular gyri and corresponds to Destrieux label S_interm_prim-Jensen. It ranks third in BCP and first in both ABCD and HCP-D.

Individual ranks differ between samples. The posterior lateral fissure ranks 30, 20, and 18 in BCP, ABCD, and HCP-D. The pericallosal sulcus ranks 51, 55, and 7. These observations do not establish developmental trajectories because the cohorts differ in acquisition and overlap in age.

**Figure S3 Regional asymmetry across cohorts**

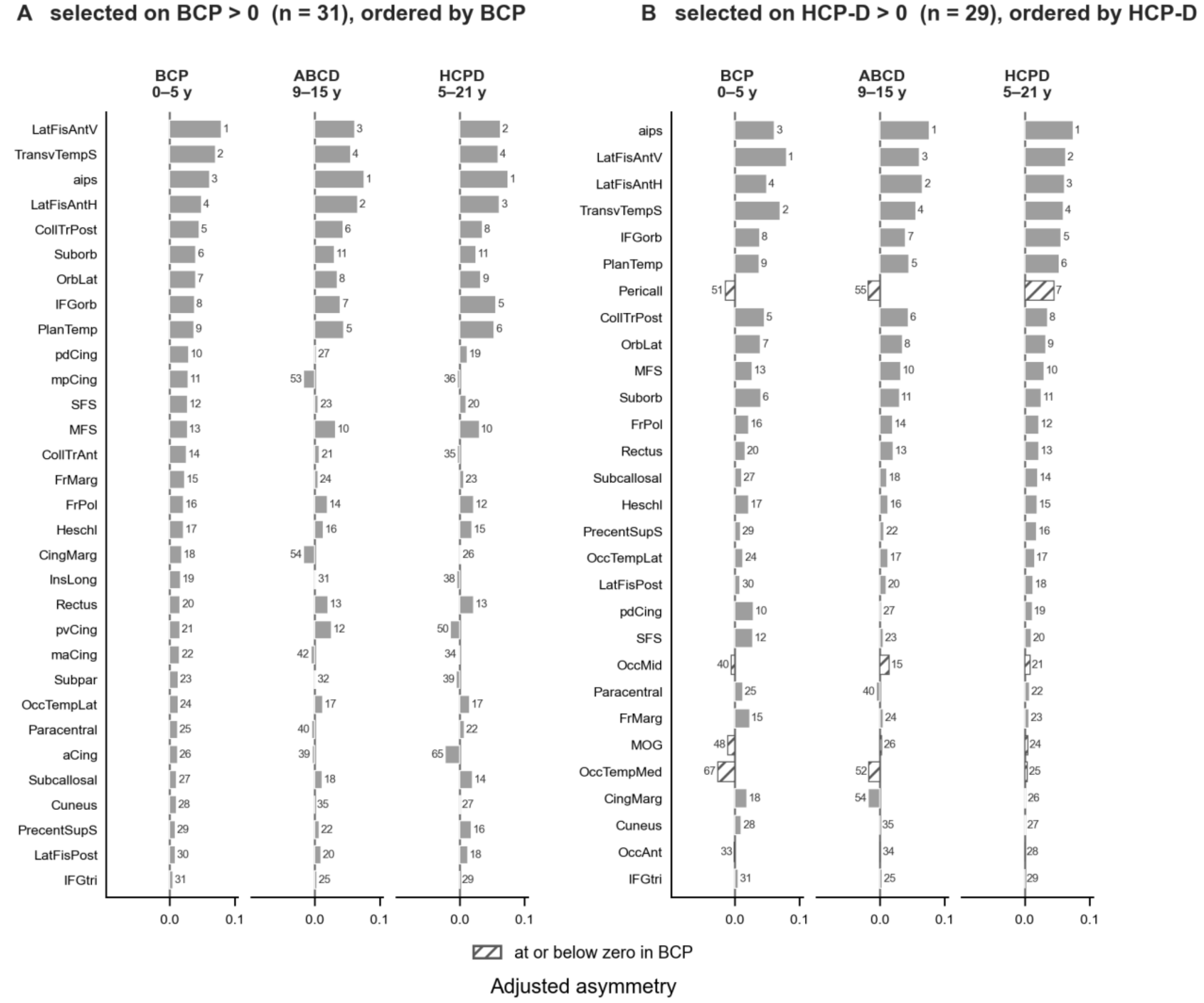


(A) The 31 regions with positive adjusted asymmetry in BCP, ordered by their BCP values. (B) The 29 regions with positive adjusted asymmetry in HCP-D, ordered by their HCP-D values. Values for all three cohorts and within-cohort ranks among 74 regions are shown for each selection. Panel B retains the ordering in main Fig. 1C to make cohort comparisons explicit. Adjusted asymmetry is the total-overlap-adjusted residual defined in section S2.

The cohort-specific overlap-adjustment slopes range over 21.6% of their mean (0.04051). Replacing them with the pooled slope nevertheless leaves regional ordering almost unchanged (Table S2). Adding family count to the adjustment reduces the HCP-D residual's correlation with family count from −0.331 to +0.004. The original and coverage-adjusted axes correlate +0.917; the mean-asymmetry–age-effect correlation changes from +0.231 to +0.221, and the interaction from +0.558 to +0.563.

**Table S2 Dependence on total overlap and stability of the adjusted axis**

| Quantity | BCP | ABCD | HCP-D |
|---|---|---|---|
| Raw asymmetry versus total overlap ρ | +0.785 | +0.822 | +0.824 |
| Cohort-specific adjustment slope | 0.04476 | 0.04076 | 0.03602 |
| Regional rank correlation after pooled-slope substitution | +0.993 | +0.99997 | +0.992 |

The mean of the cohort-specific slopes is 0.04051; their range is 21.6% of that mean. Pooled-slope substitution retains the fitting scheme and changes only the slope used for adjustment.

### S4 Connectional density and static callosal involvement

#### Degree correlations and controls

Main Fig. 2 uses cortical targets only in the outcome. The inverse relationship between adjusted asymmetry and cortical degree is present between regions and within participants or BCP sessions in all three cohorts, and remains negative under each separate control (Table S3). Tract-family count correlates with HCP-D cortical degree at +0.518 across regions (+0.484 within participants), and cortical and non-cortical degree correlate +0.632 (+0.596 within participants). The controlled correlations therefore describe different shared or residual components of the association; they are not independent effects. The tract-family control is not available for ABCD, whose consolidated tract-by-region tables retain per-family detail for five language regions only.

**Table S3 Asymmetry and cortical degree under separate controls, three cohorts**

| Additional control | HCP-D Region ρ | HCP-D Median ρ (% negative) | BCP Region ρ | BCP Median ρ (% negative) | ABCD Region ρ | ABCD Median ρ (% negative) |
|---|---|---|---|---|---|---|
| None | −0.607 | −0.300 (99.7%) | −0.720 | −0.312 (99.3%) | −0.559 | −0.302 (99.7%) |
| Total overlap | −0.611 | −0.303 (99.7%) | −0.760 | −0.324 (99.3%) | −0.559 | −0.303 (99.7%) |
| Tract coverage | −0.539 | −0.271 (99.8%) | −0.607 | −0.256 (97.5%) | — | — |
| Non-cortical degree | −0.436 | −0.219 (97.6%) | −0.546 | −0.225 (96.4%) | −0.448 | −0.232 (98.3%) |

74 homotopic regions; HCP-D, 636 participants; BCP, 446 sessions; ABCD, 1,000 participants. Within-observation values are median correlations across participants or BCP sessions; parentheses give the percentage negative. Controls are applied separately, using Spearman partial correlations. The tract-family control is unavailable for ABCD (dash). Spin $P < 0.001$ for each uncontrolled region-level correlation (10,000 rotations).

Non-cortical degree. The relationship with non-cortical degree is −0.496, −0.601, and −0.374 across regions in HCP-D, BCP, and ABCD, and −0.207, −0.220, and −0.199 as median within-observation correlations (98.3%, 97.8%, and 97.5% negative). In HCP-D, both region-level maps have split-half reliabilities above 0.998 (cortical 0.9994, non-cortical 0.9988), and disattenuation changes the controlled estimates by less than 0.003. Cortical degree given non-cortical degree is −0.436 and non-cortical given cortical is −0.182 at full target breadth (147 versus 18 targets). Recomputing cortical degree from 18 randomly chosen cortical targets, matching the non-cortical target count, gives −0.300 and −0.339 respectively: once target counts are matched the two partial correlations are similar in magnitude. This comparison indicates sensitivity to target number and does not establish stronger anatomical specificity for cortical connectivity.

Gradient. Mean cortical degree by low, middle, and high regional asymmetry tertiles has medians across participants or BCP sessions of 287, 169, and 112 streamlines per target in HCP-D; 337, 253, and 128 in BCP; and 206, 133, and 75 in ABCD. Relative to each cohort's low tertile these are 1.00 / 0.59 / 0.39, 1.00 / 0.75 / 0.38, and 1.00 / 0.64 / 0.36: the cohorts differ in absolute level but have similar relative high-tertile values. Absolute level is not compared across cohorts, because brain size, voxel geometry, and acquisition can contribute to these differences; the main figure shows the relative form.

Sign consistency. All 24 HCP-D regions in the high-asymmetry, low-degree group (above-median asymmetry and below-median degree) show sign consistency above chance after Bonferroni correction across all 332 tested family–region cells. The directional results are described in section S3. These checks do not exclude shared imaging biases.

Edge-definition sensitivity. Counting only streamlines that terminate in both regions (endpoint rule), rather than those passing through both, yields a sparser graph (18.9, 23.5, and 20.4 within-hemisphere partners per region versus 48.0, 52.7, and 41.6). The region-level correlation is −0.617, −0.596, and −0.602 (spin P = 0.001 in each cohort, 2,000 rotations; the only rotations reaching the observed value are two identity permutations, and none exceeds it), with 100.0%, 99.8%, and 99.9% of participants or BCP sessions negative, and the total-overlap control gives −0.643, −0.653, and −0.619. The inverse association is retained under both edge rules.

Streamline-length sensitivity in infancy. Because tracking is performed in native space, the 30 mm minimum streamline length is a larger fraction of an infant brain than of an adolescent one. Regenerating all BCP sessions with a 20 mm minimum changes the region-level correlation from −0.720 to −0.721; the regional degree vectors under the two settings correlate +0.9996. The regional association is insensitive to this change in minimum streamline length.

Within- and cross-hemisphere arms. Cortical degree can be split into within-hemisphere and cross-hemisphere components. In HCP-D the within arm tracks the total (−0.592 against −0.607 between regions) because it is much the larger share. The cross arm is negative under every control (between regions −0.292 raw, −0.296 given overlap, −0.468 given coverage, −0.111 given non-cortical degree; within participants −0.159, −0.165, −0.210, and −0.082, the last negative in 73% of participants). These results support lower overall connectional density, with the within-hemisphere component contributing most to total degree; they do not establish hemisphere specificity.

#### Callosal involvement

Static relationships with callosal involvement differ from the degree results (Table S4). The most asymmetric tertile has mean callosal overlap 0.155 versus 0.051 in the least asymmetric BCP tertile, approximately threefold higher. HCP-D's corresponding means are 0.114 and 0.118. These are descriptive comparisons of location, not equivalence tests.

On raw asymmetry, callosal overlap correlates negatively in BCP (−0.163; spatial P = 0.284) and HCP-D (−0.197; spatial P = 0.085), but neither relationship exceeds the spatial null. Raw asymmetry scales strongly with total association overlap, and the adjusted results do not preserve a consistent inverse callosal relationship. The tract-coverage correlation with adjusted asymmetry is −0.331 in HCP-D, compared with −0.607 for cortical degree, −0.496 for non-cortical degree, and −0.026 for callosal overlap.

**Table S4 Static relationships of asymmetry with callosal involvement**

| Quantity | BCP | ABCD | HCP-D |
|---|---|---|---|
| Adjusted asymmetry versus callosal overlap ρ | +0.298 | −0.128 | −0.026 |
| Spatial-null P for that correlation | 0.053 | — | 0.859 |
| Raw asymmetry versus callosal overlap ρ | −0.163 | — | −0.197 |
| Spatial-null P for raw-axis correlation | 0.284 | — | 0.085 |
| Mean callosal overlap in low asymmetry tertile | 0.051 | — | 0.118 |
| Mean callosal overlap in middle tertile | 0.071 | — | 0.100 |
| Mean callosal overlap in high tertile | 0.155 | — | 0.114 |

Tertiles are defined using adjusted asymmetry. Neither raw-asymmetry correlation exceeds its spatial null. A dash means not reported here, not zero or a nonsignificant test.

The descriptive asymmetry–callosal plane places perisylvian language regions at low callosal overlap and cingulate or pericallosal regions at high overlap in both displayed cohorts. The four-regime hulls in main Fig. 3 are visual guides only. Individual region assignments depend on the partition rule, and no inferential result relies on membership.

### S5 Developmental models and sensitivity analyses

#### Mean asymmetry and age effects

BCP shows a negative mean-asymmetry–age-effect relationship, including when the two quantities are estimated from disjoint participant groups (Table S5). The disjoint-subject estimate is attenuated by 14.7% relative to the whole-sample estimate. Between-region standard deviation declines by 11.5% across the sample-size-matched age bins and by 12.1% after sampling-error correction.

The corresponding between-bin dispersion difference is +0.3% in ABCD and +6.1% in HCP-D. HCP-D's positive mean-asymmetry–age-effect correlation does not exceed the spatial null despite a positive disjoint-subject estimate and is treated as unestablished. Disattenuating the three cohort correlations for map reliability yields −0.501, +0.010, and +0.280; this does not change the spatial inference.

**Table S5 Primary developmental coefficients and illustrative slopes**

| Quantity | BCP | ABCD | HCP-D |
|---|---|---|---|
| Mean-asymmetry–age-effect correlation | −0.413 | +0.009 | +0.231 |
| Spatial-null P for mean-asymmetry–age-effect correlation | 0.0022 | 0.939 | 0.053 |
| Disjoint-subject mean-asymmetry–age-effect correlation | −0.352 | — | +0.193 |
| Disjoint-subject replications in reported direction | 100% negative | — | 99% positive |
| Reliability-adjusted mean-asymmetry–age-effect correlation | −0.501 | +0.010 | +0.280 |
| Change in between-region dispersion | −12.1% | +0.3% | +6.1% |
| Standardized callosal interaction | −0.142 | +0.418 | +0.558 |
| Composite spatial-null P for interaction model | 0.336 | 0.028 | 0.0008 |
| Callosal main effect in age-effect model | −0.26 | −0.22 | +0.016 |
| Separate below-median subgroup slope | −0.356 | −0.216 | −0.062 |
| Separate above-median subgroup slope | −0.329 | +0.193 | +0.422 |
| Mean callosal overlap below median | 0.01 | 0.01 | 0.00 |
| Mean callosal overlap above median | 0.18 | 0.22 | 0.22 |
| Model-implied slope at below-median group mean | −0.27 | −0.20 | −0.05 |
| Model-implied slope at above-median group mean | −0.44 | +0.34 | +0.69 |

Model-implied slopes are those plotted in main Fig. 4 at each subgroup's own mean callosal overlap; separate subgroup slopes are fitted within each half and are different quantities. Callosal means are rounded as in the figure. HCP-D's callosal main effect is +0.016 (P = 0.88). The spatial tests are defined in section S2.

#### Interaction and callosal distribution

The standardized mean-asymmetry-by-callosal interactions are −0.142 in BCP, +0.418 in ABCD, and +0.558 in HCP-D. Their outcome-map spatial checks have empirical P values of 0.336, 0.028, and 0.0008, respectively. The null is defined in section S2 and should not be interpreted as an isolated moderation test.

Callosal overlap is heavily skewed, with cohort medians between 0.026 and 0.029 (Fig. S4). Consequently, median splits largely compare near-zero overlap with appreciable overlap. Table S5 distinguishes separate ordinary least-squares fits within these groups from slopes implied by the full interaction model at each group's mean callosal value. Only the model-implied slopes are drawn as the colored lines in main Fig. 4.

The callosal main effect in the age-effect model is −0.26 in BCP, −0.22 in ABCD, and +0.016 in HCP-D; the HCP-D main effect has P = 0.88. BCP’s mean-asymmetry–age-effect relationship remains similarly negative in the median and outer-tertile strata, with no detected moderation. Callosal involvement also shows the same direction of association with the amount of regional change across cohorts, but none of these associations exceeds its spatial null and they attenuate under magnitude and coverage control. This descriptive observation is not evidence for a general callosal main effect.

#### Specification, influence and geometry

Across specification and influence analyses, the interaction remains positive in ABCD and HCP-D (Table S6). Adding a quadratic in mean asymmetry and its callosal interaction increases rather than removes the primary interaction. A quadratic in callosal involvement is not supported in any cohort under the reported model, changes sign after leverage trimming, and does not track the ordering of the primary interactions across cohorts.

Callosal overlap correlates −0.70 to −0.75 with distance from the midline. Distance adjustment retains positive interactions, as does removal of medial-wall regions, but neither analysis uniquely separates anatomical organization from distance-related artifact. At matched ages, HCP-D's interaction is +0.325 compared with +0.418 in ABCD. Only HCP-D is restricted because the common window is ABCD's own 8.9–15.7-year range. This uses a wide age window and does not resolve the weak estimates in narrower bins.

**Table S6 Interaction sensitivity in the two older cohorts**

| Specification | ABCD | HCP-D |
|---|---|---|
| Whole-cohort primary interaction | +0.418 | +0.558 |
| Range across specification and influence checks | +0.327 to +0.640 | +0.520 to +0.659 |
| Adjusted for midline distance | +0.470 | +0.387 |
| Medial-wall regions excluded | +0.305 | +0.462 |
| Matched 8.9–15.7-year range | +0.418 | +0.325 |

Values are standardized interactions. The specification and influence range includes alternative curvature specifications and influence trims; the trims remove regions from the linear primary model as defined in section S2. Only HCP-D is age-restricted in the matched comparison; ABCD retains its full 8.9–15.7-year range.

**Figure S4 Distribution of regional callosal overlap**

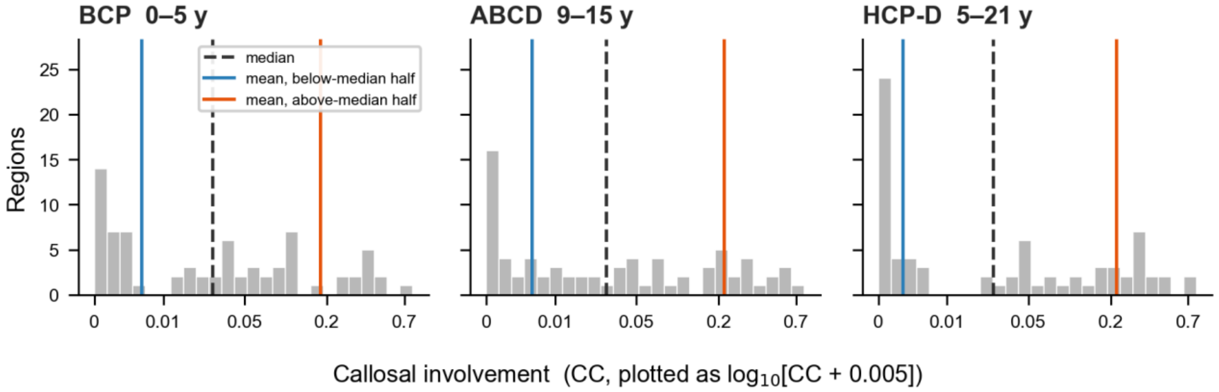


Distribution of cohort-mean callosal overlap across the 74 regions in each cohort. The dashed black line marks the median; colored lines mark the means of the below- and above-median groups. The horizontal coordinate is $\log_{10}$(callosal overlap + 0.005), with tick labels giving overlap values. The skew explains why a median split separates near-zero overlap from appreciable overlap and why the two group means differ substantially from the median.

Specification, influence, geometry, and disjoint-participant results are summarized in the text and Tables S5–S6.

### S6 Longitudinal estimates and age windows

#### Within-participant change and regional inference

ABCD's primary longitudinal analysis pools wave-adjacent nominal two-year transitions: baseline to two years (6,281 pairs) and two to four years (2,231 pairs), totaling 8,512 pairs. Its standardized interaction is +0.316, with 73.8% of the pooling weight from baseline to two years. The mean per-wave cross-sectional interaction is +0.311. A sensitivity pooling all ordered visit pairs, including four-year intervals, yields +0.286 over 10,868 pairs (Table S7). Counts refer to visit pairs, not independent participants.

For the primary 8,512-pair analysis, the ordinary least-squares P value is 0.044, compared with 0.006 under precision weighting and 0.092 under the spatial null; the bias-corrected and accelerated (BCa) bootstrap interval is [−0.023, +0.806] (4,000 region resamples with jackknife acceleration). Precision weights correlate −0.505 with mean regional asymmetry. Adding its quadratic reduces the gap between weighted and unweighted estimates from 41.9% to 11.5%. The convergence with cross-sectional estimates is therefore one of direction and magnitude, with estimator-sensitive inference.

The anchor sensitivity instead pairs each participant's visit with the next available visit. Its 8,917 pairs include 405 baseline-to-four-year pairs from participants without a two-year visit; these account exactly for the difference from the primary sample. Both analyses draw from the joint pool of 19,569 sessions from 10,652 participants. The anchor interaction is +0.3073 (ordinary least-squares P = 0.051; HC3 P = 0.215). Its 95% percentile region-bootstrap interval is [+0.008, +0.898], from 4,000 resamples of the 74 regions with replacement. A separate pair-bootstrap interval is [+0.211, +0.389], from 2,000 pair resamples (seed 31). Pair resampling does not preserve participant clusters; region resampling does not preserve spatial dependence. Neither interval replaces the primary sample's BCa interval or spatial test.

**Table S7 ABCD longitudinal estimates and uncertainty**

| Analysis and quantity | Estimate or count |
|---|---|
| Primary wave-adjacent two-year analysis | |
| Standardized interaction | +0.316 |
| Consecutive visit pairs | 8,512 |
| Baseline-to-two-year pooling weight | 73.8% |
| Ordinary least-squares P | 0.044 |
| Precision-weighted P | 0.006 |
| Spatial-null P | 0.092 |
| BCa bootstrap interval | [−0.023, +0.806] |
| **Other primary-analysis summaries** | |
| All ordered pairs interaction / pair count | +0.286 / 10,868 |
| Mean per-wave cross-sectional interaction | +0.311 |
| Anchor next-available-visit fit | |
| Standardized interaction / pair count | +0.3073 / 8,917 |
| Ordinary least-squares P | 0.051 |
| HC3 P | 0.215 |
| Region-bootstrap 95% percentile interval | [+0.008, +0.898] |
| Individual / region-mean change variance ratio | 628 |
| **Visit transitions** | |
| Baseline to two years interaction | +0.371 |
| Baseline to four years interaction | +0.228 |
| Two to four years interaction | +0.107 |

The cohort-mean regional asymmetry is the predictor; the outcome summarizes within-participant change by region. Primary wave-adjacent pairs and next-available-visit anchor pairs are distinct. The anchor region-bootstrap interval is a 95% percentile interval; it is not BCa or a spatial-null interval.

#### Differences between transitions

Transition-specific interactions are +0.371 for baseline to two years, +0.228 for baseline to four years, and +0.107 for two to four years. Their ordering does not follow a simple interval-width-only attenuation prediction, under which the four-year estimate would exceed the two-year estimates. These are descriptive comparisons, not evidence for a monotonic change in moderation with age.

The two nominal two-year transitions share 1,951 participants with all three waves, including the middle visit with opposite roles in the two changes. Their coefficient estimates are therefore dependent. The transition ordering is interpreted descriptively, without a formal test of between-transition differences.

#### Cross-sectional age window width

At fixed subsample size of 100, the HCP-D interaction increases from a median +0.107 over three-year windows to +0.390 over sixteen-year windows (Fig. S5). These results are compatible with attenuation in narrow windows, but expanding a window also changes the ages included. Window width and developmental age remain confounded. This analysis concerns cross-sectional estimation windows and should not be applied directly to within-person follow-up intervals.

#### Individual change and measurement noise

For participants with all three waves, the shared-wave model writes the two changes as $\Delta_{01} = \tau + e_1 - e_0$ and $\Delta_{12} = \tau + e_2 - e_1$. Here τ is a persistent trajectory component and the e terms are independent measurement errors, independent of τ and equal in variance across waves. With $k = \mathrm{Var}(\tau)/\mathrm{Var}(e)$, the predicted correlation is $r = (k - 1)/(k + 2)$; pure measurement error gives $r = -0.5$. Across participant-by-region cells from 1,951 participants, the observed r is −0.4613, with a participant-bootstrap standard error of 0.0048 (1,000 resamples). Inverting the model gives $k = (1 + 2r)/(1 - r) = 0.0529$ and a trajectory share $k/(k + 2) = 2.58\%$ of change variance. The complementary 97.4% is assigned to measurement error under these assumptions, rather than measured directly. Unequal error variances or interval-specific true changes would alter the decomposition.

In the 8,917-pair anchor analysis, individual-change variance is 628 times the between-region variance of mean change. Mean absolute individual change is 0.0232 and mean absolute region-mean change is 0.0011, approximately 93% and 4% of the between-region mean-asymmetry standard deviation (0.0249). The corresponding primary-sample absolute changes are 0.0231 and 0.0010. These scales characterize different units and are not individual-level prediction accuracy.

**Figure S5 Cross-sectional age window width**

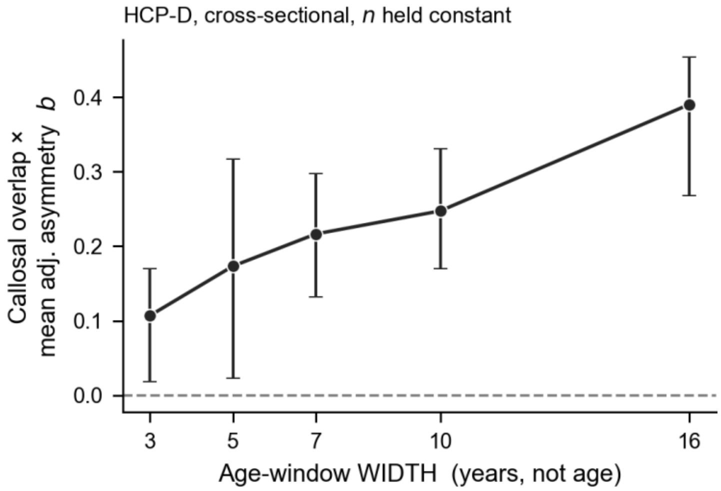


HCP-D standardized callosal interaction across age windows of 3, 5, 7, 10, and 16 years, holding each subsample at 100 participants. Markers show median estimates across resamples and bars the interquartile range, not confidence intervals. Median interaction increases from +0.107 for a three-year window to +0.390 for a sixteen-year window. Changing window width also changes the age span sampled.

### S7 Map reliability and anatomical change

#### Reliability and map correspondence

Mean-asymmetry maps have split-half reliability of approximately 0.99. Age-effect maps are less reliable, especially in BCP and HCP-D (Table S8). Correction for reliability therefore mainly addresses attenuation of the change maps.

Within ABCD, consecutive-interval region-mean change maps have Pearson r = +0.721 (Spearman ρ = +0.739), and the cross-sectional and within-subject maps correlate +0.943. Comparisons share participants and scanners; consecutive intervals share the middle wave. Across cohorts, observed

ABCD–HCP-D agreement is +0.53, with a disattenuated correlation of +0.657 (reported interval [+0.414, +0.861]). The within-subject ABCD comparison with HCP-D is +0.479 observed and +0.603 after disattenuation. The BCP–HCP-D interval [−0.487, +0.202] does not overlap the ABCD–HCP-D interval, whereas the BCP–ABCD interval [−0.116, +0.461] does. Interval overlap is descriptive, not a formal difference test; comparisons do not isolate age from cohort or acquisition effects.

BCP's regional convergence is supported by disjoint-participant estimation (ρ = −0.352; all 200 replications negative) and by the reliability-adjusted mean-asymmetry–age-effect association (−0.501 versus −0.413 before adjustment). These controls address different questions from a comparison of age-bin maps.

**Table S8 Map reliability and cross-map correspondence**

| Map or comparison | Value | Statistic |
| --- | --- | --- |
| BCP age-effect map | 0.675 | Corrected split-half reliability |
| ABCD cross-sectional age-effect map | 0.947 | Corrected split-half reliability |
| ABCD within-subject change map | 0.916 | Corrected split-half reliability |
| HCP-D age-effect map | 0.691 | Corrected split-half reliability |
| Mean-asymmetry maps | ≈0.99 | Corrected split-half reliability |
| ABCD consecutive-interval maps | +0.721 | Observed Pearson r |
| ABCD cross-sectional versus within-subject map | +0.943 | Observed correlation |
| ABCD cross-sectional versus HCP-D | +0.53 | Observed correlation |
| ABCD cross-sectional versus HCP-D | +0.657 | Disattenuated correlation |
| ABCD within-subject versus HCP-D | +0.479 | Observed |
| ABCD within-subject versus HCP-D | +0.603 | Disattenuated correlation |
| BCP versus HCP-D | −0.10 | Observed correlation |
| BCP versus HCP-D | −0.148 | Disattenuated correlation |
| BCP versus ABCD | +0.169 | Disattenuated correlation |
| ABCD cross-sectional versus HCP-D interval | [+0.414, +0.861] | Disattenuated |
| BCP versus HCP-D interval | [−0.487, +0.202] | Disattenuated |
| BCP versus ABCD interval | [−0.116, +0.461] | Disattenuated |

Whole-map split-half reliabilities are Spearman–Brown corrected. Map correlations are descriptive, with no spatial-null test. ABCD comparisons within the dataset share participants, and consecutive intervals share the middle wave. The consecutive-interval Pearson and Spearman statistics are distinguished explicitly.

### Shared regional changes

The eight-region set in main Figs. 5–6 is the intersection of ABCD's and HCP-D's 15 largest absolute regional age effects: gyrus rectus, precuneus, subcallosal gyrus, H-shaped orbital sulcus, parahippocampal gyrus, orbital gyrus, parieto-occipital sulcus, and dorsal posterior cingulate gyrus. Their median distance from the midline is 11.2 mm, compared with 35.1 mm for all 74 regions.

Re-estimating ABCD age effects using only follow-up waves retains seven of the eight regions; precuneus is replaced by transverse temporal sulcus. This sensitivity separates the baseline wave from the waves used to estimate change, but does not by itself make all regional quantities statistically independent. Main Fig. 5 displays 16 FDR-significant regional age effects in BCP, 47 in ABCD, and 13 in HCP-D. Because significance is calculated across sessions, it may be anti-conservative where participants contribute repeated scans. The shared set is descriptive anatomical convergence, not an additional inferential test of replication.

### Tract attribution

#### Figure S6 Tract contributions to regional age effects

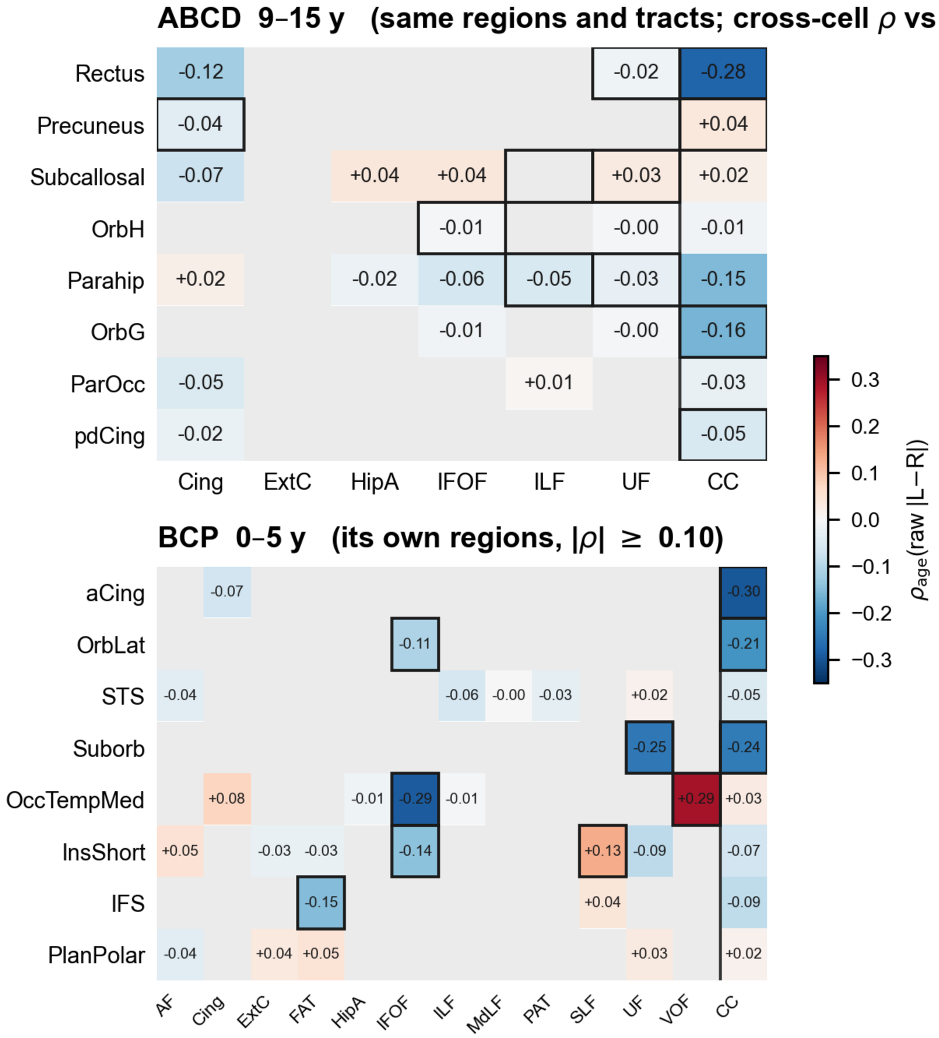


Top: ABCD tract-by-region age effects for the eight-region set and tract families displayed for HCP-D in main Fig. 6. The displayed cross-cell correlation is +0.60 across 24 cells, including the callosal column. Bottom: BCP effects for its own selected regions, with an absolute age-correlation display threshold of 0.10. Association-family cells show age correlations of raw absolute left–right differences; the CC column shows age correlations of callosal overlap. Gray cells fail the 50% coverage criterion. Outlines identify rendered examples. These heatmaps describe anatomical contributions and do not retest the regional interaction.

Tract-by-region age effects use raw absolute left–right differences because overlap adjustment is defined at regional level. Family cells require at least 50% coverage; the callosal column in main Fig. 6 instead gives the age effect on callosal overlap. Across 24 shared cells including this column, ABCD and HCP-D have Spearman ρ = +0.601 (nominal P = 0.0019; Fig. S6). Restricting the comparison to the 16 shared association-family cells gives ρ = +0.668 (nominal P = 0.005). BCP–HCP-D correspondence is +0.059 over 26 available comparison cells. Across the 16 shared family cells, median absolute age correlation is 0.0216 in ABCD and 0.0633 in HCP-D, a ratio of 0.34. These are descriptive cell comparisons; nominal P values do not account for spatial or cell dependence.

BCP renderings use its own large age effects, with absolute correlations ≥0.10, because its largest changes do not concentrate in the older-cohort shared set. Raw asymmetry decreases at most of these displayed BCP pairs. One callosal projection to lateral orbital cortex lacks template streamlines even after four-voxel dilation and is not rendered. Bundle-color saturation is floored for visibility; exact magnitudes are given in the heatmap, not by visual saturation.

### S8 Sex comparisons

Mean-asymmetry and callosal maps are highly similar between sexes in all three cohorts (Table S9), with no detected difference in mean magnitude. The static asymmetry–callosal relationship has the same sign in both sexes within each cohort. The interaction is positive in both sexes in ABCD and HCP-D, and the male-minus-female difference interval includes zero in all three cohorts.

Male–female age-effect-map correlations are similar to random-half correlations calculated at comparable sample sizes. These random-half values differ from the Spearman–Brown-corrected whole-map reliabilities in Table S8 and should not be interchanged. One descriptive exception is the already unestablished mean-asymmetry–age-effect association in HCP-D: the sex-stratified value is +0.298 in females and +0.015 in males. These values do not establish a sex difference in the primary interaction.

**Table S9 Sex-stratified maps and interaction estimates**

| Quantity | BCP | ABCD | HCP-D |
|---|---|---|---|
| Male–female mean-asymmetry map ρ | +0.977 | +0.991 | +0.979 |
| Male–female callosal-overlap map ρ | +0.971 | +0.997 | +0.991 |
| Male–female age-effect-map ρ | +0.581 | +0.885 | +0.519 |
| Comparable random-half age-map correlation | 0.509 | 0.899 | 0.528 |
| Interaction in males / females | −0.235 / −0.067 | +0.423 / +0.396 | +0.456 / +0.522 |
| Male-minus-female difference and reported interval | −0.168 [−0.424, +0.139] | +0.027 [−0.093, +0.144] | −0.066 [−0.261, +0.152] |

Interaction coefficients are labeled by sex; differences are male minus female. Difference intervals are 95% percentile intervals from 200 resamples within each sex, using sessions as the resampling unit (seed 2027). This does not preserve participant clustering in BCP or ABCD; the intervals are descriptive and are not equivalence tests. No interval excludes zero. BCP has 435 sessions with a sex record (213 male, 222 female), of which 429 have usable age; ABCD has 19,569 (10,237 male, 9,332 female); HCP-D has 636 (292 male, 344 female). Random-half correlations are distinct from corrected whole-map reliabilities in Table S8.

### S9 Regional maturational change

For each region, an age-association index was computed as the Spearman correlation across sessions between age and mean bilateral association overlap: the mean of L + R across covered association families. The primary index is the absolute correlation; the signed correlation is a sensitivity. A secondary HCP-D index substitutes cortical degree for overlap. Each index enters the regional model as a covariate and as a moderator of mean asymmetry, using the same vector shown in Fig. S7. None estimates maturational onset: both a region whose change has already occurred and one whose change has not begun can have a near-zero age association. Regional profiles of the index by anatomical class are shown in Fig. S8.

The primary absolute index has little association with callosal overlap in HCP-D (ρ = −0.07) or ABCD (+0.12). The signed index in ABCD is an exception (+0.26); the HCP-D index derived from the regional age correlation with cortical degree correlates +0.21 with callosal overlap. Thus the broader set of indices cannot be described as unrelated to callosal overlap in every cohort. Including the absolute index as both a covariate and a moderator of mean asymmetry leaves the callosal interaction close to its primary value: +0.537 (P = 0.0002) versus +0.558 in HCP-D, and +0.400 (P = 0.004) versus +0.418 in ABCD. Using the primary streamline-count matrices, the HCP-D degree-based sensitivity retains a positive callosal interaction of +0.512 (95% interval [+0.251, +0.773], P = 0.0002), modestly attenuated from +0.558. The degree age-association index's association with callosal overlap is sensitive to graph definition and is not interpreted as a stable directional relationship. In ABCD, a within-subject index agrees with the cross-sectional index at ρ = +0.957 and yields a callosal interaction of +0.319 (P = 0.015). These analyses address one measured aspect of maturation and do not rule out common developmental regulation of callosal connectivity and asymmetry.

**Figure S7 Regional maturational change and the callosal interaction**

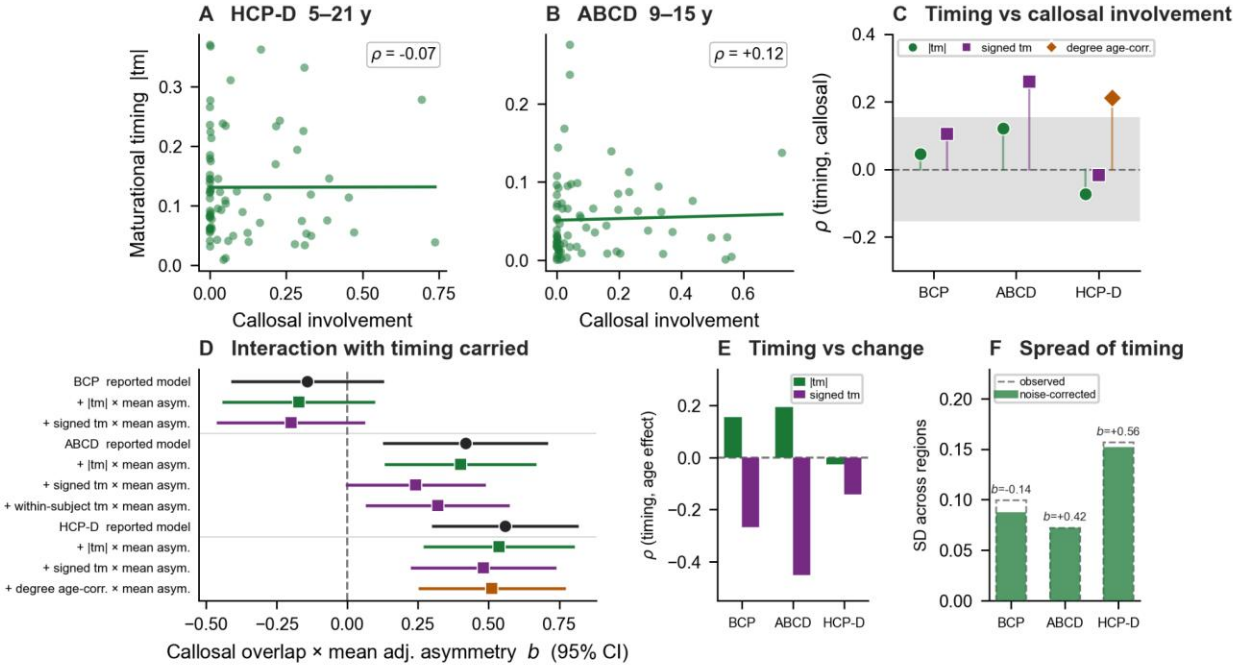


(A, B) Absolute Spearman correlation between age and mean bilateral association overlap versus callosal overlap in HCP-D and ABCD. (C) Associations between callosal overlap and the alternative indices across cohorts; the shaded |ρ| < 0.15 band is a descriptive reference, not a confidence interval or a significance threshold. The HCP-D degree age-association index uses the primary streamline-count matrices (ρ = +0.21); its association with callosal overlap is sensitive to graph definition. (D) Standardized callosal interactions before and after carrying the index as a covariate and moderator; bars are ordinary least-squares 95% t confidence intervals for the standardized callosal × mean-asymmetry coefficient, with 70 residual degrees of freedom for the four-parameter reported model and 68 for each six-parameter augmented model (index main effect and index × mean asymmetry added). The HCP-D degree age-association sensitivity is +0.512 [+0.251, +0.773] (P = 0.0002). (E) Absolute and signed indices versus the asymmetry age effect; this relationship is partly tied to the construction of the measures and is not independent validation. (F) Observed and sampling-error-corrected between-region spread of the absolute index.

**Figure S8 Regional profiles of the age association index**

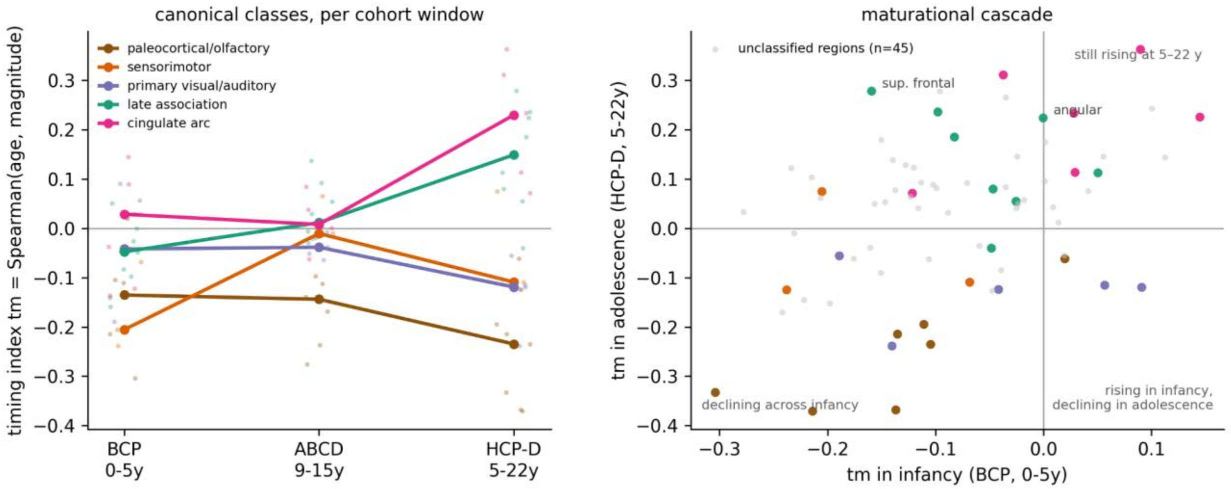


Left: regional age-association indices grouped by anatomical class within each cohort. Large connected markers summarize classes; smaller points show regions. Connecting lines compare overlapping cohort windows and do not trace within-person trajectories. Right: signed Spearman correlations between age and mean bilateral association overlap (labeled “magnitude” in the plot) for BCP and HCP-D, with selected anatomical classes highlighted. These are descriptive profiles, not independent validation or estimates of when maturation begins or peaks.

### S10 Figure abbreviations and analysis samples

Table S10 defines the tract abbreviations from the main figure legends and selected regional labels.

**Table S10 Tract and selected regional abbreviations in the main figures**

| Label | Meaning |
|---|---|
| AF | Arcuate fasciculus |
| CB; Cing | Cingulum bundle |
| ExtC | Extreme capsule |
| FAT | Frontal aslant tract |
| HipA | Hippocampal alveus |
| IFOF | Inferior fronto-occipital fasciculus |
| ILF | Inferior longitudinal fasciculus |
| MdLF | Middle longitudinal fasciculus |
| PAT | Parietal aslant tract |
| SLF | Superior longitudinal fasciculus |
| UF | Uncinate fasciculus |
| VOF | Vertical occipital fasciculus |
| CC | Corpus callosum or callosal overlap, according to the panel |
| aips | Anterior intermediate parietal sulcus; sulcus intermedius primus of Jensen; Destrieux S_interm_prim-Jensen |

| Label | Meaning |
|---|---|
| PlanTemp | Planum temporale |
| Rectus | Gyrus rectus |
| Subcallosal | Subcallosal gyrus |
| OrbH | H-shaped orbital sulcus |
| Parahip | Parahippocampal gyrus |
| OrbG | Orbital gyrus |
| ParOcc | Parieto-occipital sulcus |
| pdCing | Dorsal posterior cingulate gyrus |

Main Fig. 6 and Fig. S6 use ExtC for extreme capsule and HipA for hippocampal alveus.

Regional summaries use the sample appropriate to each analysis. Mean adjusted asymmetry, callosal overlap, age effects, and the overlap age-association index in the interaction models use the age-effect sample (HCP-D, 636 participants; BCP, 429 sessions; ABCD, 19,569 sessions). Raw asymmetry, bilateral association overlap, and family coverage were also summarized in the broader level samples (636, 450, and 19,605 observations, respectively); the last is the ABCD asymmetry-only sample, distinct from the joint sample in Table S1. Connectional-density correlations pair degree with mean adjusted asymmetry from the same graph sample (HCP-D, 636 participants; BCP, 446 sessions; ABCD, 1,000 participants).

### References


1. B. R. Howell et al., The UNC/UMN Baby Connectome Project (BCP): An overview of the study design and protocol development. NeuroImage 185, 891–905 (2019). https://doi.org/10.1016/j.neuroimage.2018.03.049
2. B. J. Casey et al., The Adolescent Brain Cognitive Development (ABCD) study: Imaging acquisition across 21 sites. Developmental Cognitive Neuroscience 32, 43–54 (2018). https://doi.org/10.1016/j.dcn.2018.03.001
3. L. H. Somerville et al., The Lifespan Human Connectome Project in Development: A large-scale study of brain connectivity development in 5–21 year olds. NeuroImage 183, 456–468 (2018). https://doi.org/10.1016/j.neuroimage.2018.08.050
4. M. P. Harms et al., Extending the Human Connectome Project across ages: Imaging protocols for the Lifespan Development and Aging projects. NeuroImage 183, 972–984 (2018). https://doi.org/10.1016/j.neuroimage.2018.09.060
5. J.-P. Fortin et al., Harmonization of multi-site diffusion tensor imaging data. NeuroImage 161, 149–170 (2017). https://doi.org/10.1016/j.neuroimage.2017.08.047
6. F.-C. Yeh, DSI Studio: An integrated tractography platform and fiber data hub for accelerating brain research. Nature Methods 22, 1617–1619 (2025). https://doi.org/10.1038/s41592-025-02762-8
7. F.-C. Yeh, V. J. Wedeen, W.-Y. I. Tseng, Generalized q-sampling imaging. IEEE Transactions on Medical Imaging 29, 1626–1635 (2010). https://doi.org/10.1109/TMI.2010.2045126
8. F.-C. Yeh, Population-based tract-to-region connectome of the human brain and its hierarchical topology. Nature Communications 13, 4933 (2022). https://doi.org/10.1038/s41467-022-32595-4
9. C. Destrieux, B. Fischl, A. Dale, E. Halgren, Automatic parcellation of human cortical gyri and sulci using standard anatomical nomenclature. NeuroImage 53, 1–15 (2010). https://doi.org/10.1016/j.neuroimage.2010.06.010
10. A. F. Alexander-Bloch et al., On testing for spatial correspondence between maps of human brain structure and function. NeuroImage 178, 540–551 (2018). https://doi.org/10.1016/j.neuroimage.2018.05.070
11. M. Catani et al., Symmetries in human brain language pathways correlate with verbal recall. Proceedings of the National Academy of Sciences USA 104, 17163–17168 (2007). https://doi.org/10.1073/pnas.0702116104
12. M. Thiebaut de Schotten et al., A lateralized brain network for visuospatial attention. Nature Neuroscience 14, 1245–1246 (2011). https://doi.org/10.1038/nn.2905
